# How relevant is the prior? Bayesian causal inference for dynamic perception in volatile environments

**DOI:** 10.1101/2024.10.29.620874

**Authors:** David Meijer, Roberto Barumerli, Robert Baumgartner

## Abstract

Interpreting sensory prediction errors can be challenging in volatile environments because they can be caused by stochastic noise or by outdated predictions. Noisy signals should be integrated with prior beliefs to improve precision, but the two should be segregated when environmental changes render prior beliefs irrelevant. Bayesian causal inference provides a statistically optimal solution to deal with uncertainty about the causes of prediction errors. However, the method quickly becomes memory intensive and computationally intractable when applied sequentially.

Here, we systematically evaluate the predictive performance of Bayesian causal inference for perceptual decisions in a spatial prediction task based on noisy audiovisual sequences with occasional changepoints. We elucidate the simplifying assumptions of a previously proposed reduced Bayesian observer model, and we compare it to an extensive set of models based on alternative simplification strategies.

Model-free analyses revealed the hallmarks of Bayesian causal inference: participants seem to have integrated sensory evidence with prior beliefs to improve accuracy when prediction errors were small, but prior influence decreased gradually as prediction errors increased, signalling probable irrelevance of the priors due to changepoints. Model comparison results indicated that participants computed probability-weighted averages over the causal options (noise or changepoint), akin to the reduced Bayesian observer model. However, participants’ reliance on prior beliefs was systematically smaller than expected, and this was best explained by individually fitting lower-than-optimal parameters for the a-priori probability of prior relevance.

We conclude that perceptual decision makers utilize priors flexibly to the extent that they are deemed relevant, though also conservatively with a lower tendency to bind than ideal observers. Simplified consecutive Bayesian causal inference predicts key characteristics of belief updating in changepoint environments and forms a suitable foundation for modelling dynamic perception in a changing world.

## 1. Introduction

Perception is inherently uncertain. Sensory signals are disturbed by noise from external sources and our nervous system suffers from imperfect processing and transmission (Faisal et al., 2008). Besides, the limited information that is conveyed by the sensory signals is often insufficient by itself to obtain an unambiguous understanding of our environment. One efficient way to nevertheless form a coherent model of the world around us is to supplement and compare incoming sensory information with expectations based on our current beliefs (Friston, 2010; Clark, 2013). A mismatch between the two generates a prediction error that calls for an explanation. Prediction errors can be attributed to sensory noise or imprecise predictions, allowing the novel sensory information to be incorporated into the pre-existing beliefs to improve their accuracy. Alternatively, prediction errors can also be caused by previously unknown or new sources of the sensory signals in a changing environment, necessitating an update to the observer’s model of the world.

Bayesian inference is a probabilistic approach to deal with perceptual uncertainty in a manner that has the potential to be statistically optimal (Ma, 2012). Accordingly, it has been proposed that our brains attenuate the detrimental impact of random noise, thus improving perceptual precision, by integrating redundant information across sensory cues and prior beliefs by means of reliability-weighted averaging (Knill & Pouget, 2004). Indeed, there is an abundance of reports that collectively demonstrate that what we perceive is often biased by our expectations and by concurrent sensory signals in accordance with Bayesian integration (de Lange et al., 2018, Meijer & Noppeney, 2020) and that our brains are in principle capable of such probabilistic computations (Pouget et al., 2013).

However, one should not integrate sensory information that does not originate from the same real-world object or event (i.e., cause). Within the Bayesian inference framework, one can naturally account for the possibility of multiple sensory sources by a hierarchical extension of the prior (Yuille & Bülthoff, 1996; Shams & Beierholm, 2010). For example, in an audiovisual ventriloquism paradigm, an observer must entertain two competing causal structures: either the auditory and visual signals stem from the same source location, or they come from two different locations (Körding et al., 2007). Moderately small spatial disparities between the sensory signals could have easily been caused by noise, so an observer may falsely infer a common source and thus integrates the two sensory signals, leading to the ventriloquist illusion. Yet, the illusion does not occur for larger spatial disparities when it is more likely that the two signals originated from two different locations. Results from a multitude of psychophysics and neuroimaging studies support the claim that our brains apply such Bayesian causal inference to determine whether to integrate or segregate sensory signals (Shams & Beierholm, 2022).

Another field of research in which Bayesian inference has been successful at describing human perceptual decisions is concerned with learning about changing environments (Behrens et al., 2007; O’Reilly, 2013; Kang et al., 2024). Of specific interest to us here is the problem of changepoint detection: i.e., how does one determine whether their current model of the world has become irrelevant because of a sudden change in their surroundings? In other words, when should one integrate noisy sensory evidence with previously held beliefs to improve precision, and when should one segregate the two because the latest sensory signal stems from a novel reality? The problem is thus essentially one of causal inference (see Figure 1) and Bayesian probability theory prescribes the optimal approach to dynamically deal with the combined uncertainty from stochastic noise and potential changepoints in the generative process (Adams & MacKay, 2007; Fearnhead and Liu, 2007).

**Figure 1.**
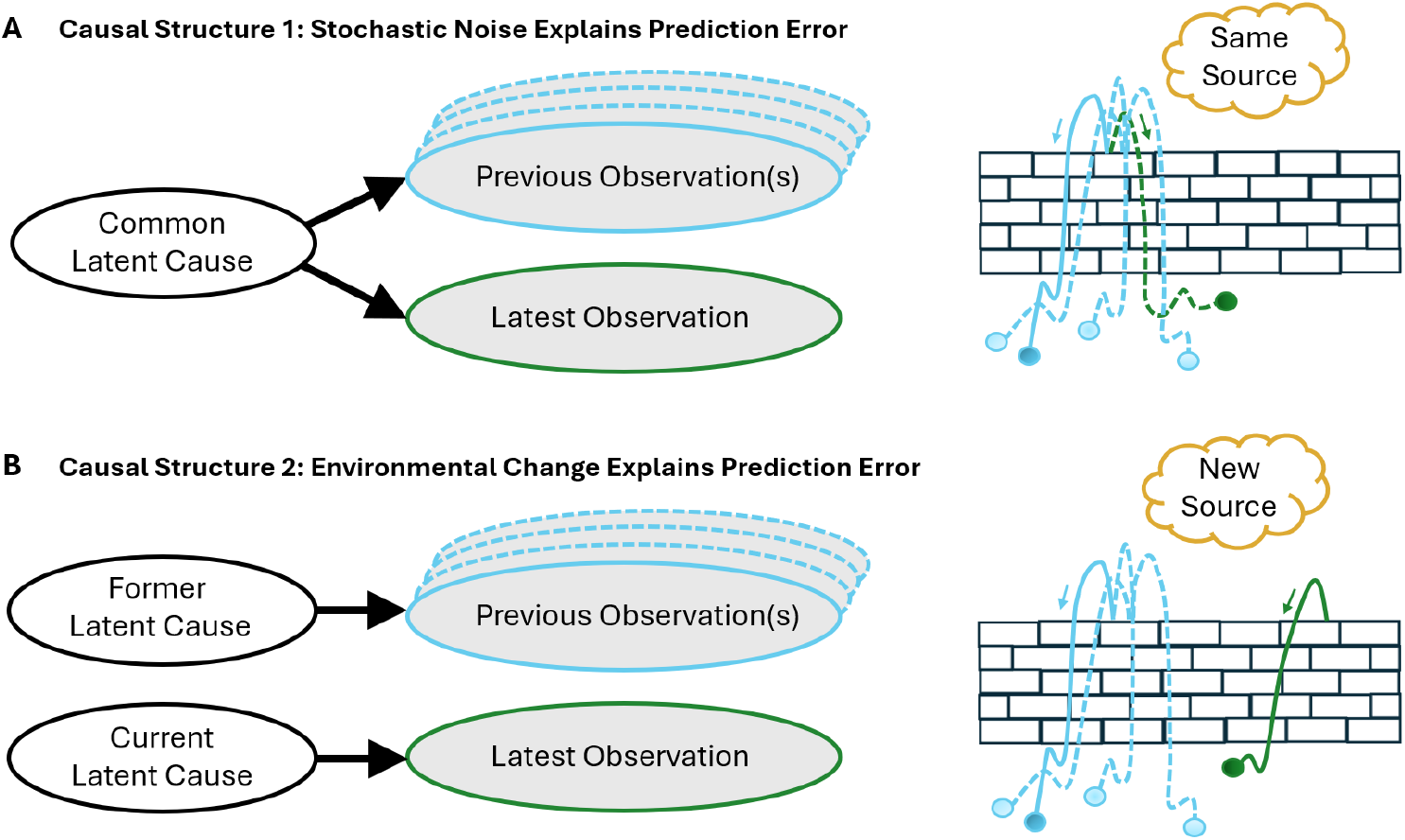
**Two competing models of the world for explaining prediction errors in a changepoint paradigm. A). The first causal structure assumes that the latest observation stems from the same source as the preceding observations. B). The second causal structure instead assumes that the latest observation originates from a different source. Note that the true causes that give rise to the sensory signals are unknown to observers (i.e., latent). This is illustrated on the right side by a brick wall that obscures the true source location(s) of the balls that are thrown over it (i.e., noisy observations). The inferred cause for the latest observation is mentioned in the thought clouds.**

The fully Bayesian solution to the changepoint detection problem allows an agent to retrospectively evaluate evidence about changepoints that occurred several observations in the past. While such inference with hindsight leads to the most precise beliefs about the current state of the world, it also requires extensive memory and computational capacity that seems implausible for the human brain. Hence, simplified near-Bayesian changepoint models were developed that approximate the optimal solution and provide more realistic algorithms that may be implemented by the brain (Nassar et al., 2010, 2012; Wilson et al., 2013). Indeed, available psychophysical, neuroimaging and neurophysiological work in combination with computational modelling generally support the hypothesis that the brain relies on approximately-Bayesian inference for adaptive learning in changing environments (Yu et al., 2021).

Here, our main contributions are threefold. First, we attempt to bridge a gap between the fields of learning and sensory cue integration by presenting a previously proposed reduced Bayesian observer model for changepoint detection (Nassar et al., 2012) in a way that is readily understandable by those familiar with the Bayesian causal inference models for sensory cue integration. In so doing, we subtly but fundamentally shift the perspective from the updating of beliefs when new information becomes available (e.g., learning rates), to the interpretation of sensory signals by latent causal inference (Gershman et al., 2015). While the mathematical foundation of these modelling perspectives is identical in the changepoint detection paradigm, we argue that Bayesian causal inference provides a flexible normative framework that lends itself well to model extensions and generalizations across tasks (Shams & Beierholm, 2022). By elucidating its core components for consecutive (as opposed to concurrent) cue integration, we lay-out key predictions and introduce testable hypotheses for approximate Bayesian causal inference in the domain of dynamic belief updating.

Second, we evaluate the reduced Bayesian observer model’s performance in explaining spatial predictions of participants in a changepoint detection task wherein expectations had to be formed and updated during the ongoing presentation of stimulus sequences. Participants were prompted for a response at unpredictable times when the sequence suddenly stopped. This experimental design is in stark contrast to the more commonly used procedure where participants are required to make a prediction response after every stimulus (Nassar et al., 2010, 2012, 2019, McGuire et al., 2014). Such sequence disruptions likely affect consecutive cue integration efficacy. Instead, by using a task design that requires covert evidence accumulation rather than overt serial decision-making, we aim to examine Bayesian causal inference as a mechanism of implicit sequential perception, while also aspiring to avoid explicit choice history biases (Talluri et al., 2018; Bévalot & Meyniel, 2023). We adopt the rich behavioral dataset that was previously analysed by Krishnamurthy, Nassar, Sarode and Gold (2017), but we limit our analysis to the audiovisual sequences and the prediction responses, whereas their original publication focused on pupillometry recordings and localization responses about sounds that followed those predictions.

Third, we derive the fully Bayesian solution for this experimental paradigm and subsequently systematically introduce simplifications and alternative modelling assumptions. This allows us to build a factorial framework of near-Bayesian changepoint detection model variations with varying complexity and decisional strategies. We then fit all models to the behavioral data and perform a model comparison to evaluate which of the tested algorithmic components are most probably utilized by the brain. As such, we provide a systematic evaluation of memory and complexity reduction in sequential Bayesian causal inference models for dynamic perception in volatile environments. Please note that we restrict our analyses to modelling static sources of noisy sensory signals with sudden environmental changes. In the discussion section, we allude to possible model extensions that allow the tracking of dynamically moving sources in addition to changepoints (Nassar et al., 2010, Nassar 2019), and we compare Bayesian causal inference against alternative approaches that model environmental volatility through moving sources exclusively (i.e., hierarchical extensions of the Kalman filter; Mathys et al. 2011, 2014; Piray & Daw, 2020). Finally, we highlight possible avenues for future research that can expand sequential Bayesian causal inference models to incorporate latent causes with higher order statistical regularities (Skerritt-Davis & Elhilali, 2018, 2021) or for use in environments with multiple sources that are simultaneously active (Gershman et al., 2015; Larigaldie et al., 2024).

## 2. Results

Twenty-nine participants performed a spatial prediction task in which they indicated the anticipated location of an upcoming stimulus by means of a mouse response on a semi-circular arc after having been presented with a sequence of audiovisual stimuli (Figure 2A). Responses were self-paced, and the median reaction time was 1.72 (IQR: 1.49–2.35) seconds. The sequences mostly allowed for reasonable predictions because the stimuli locations (at times *t*), *x*_*t*_, were sampled from a normal distribution, centred on the ‘generative mean’ *μ*_*t*_ and with width defined by the ‘experimental noise’ σ_*exp*_ (SD = 10° or 20° in blocked low or high noise conditions, respectively). Hence, the mean location of the preceding stimuli provides a good prediction for the location of the next stimulus. However, at every timepoint *t*, there was a 15% chance for a changepoint (hazard rate *H*_*cp*_ = 0.15), in which case the generative mean *μ*_*t*_ was resampled at random from within the space boundaries (uniform distribution from *a* = −90° to *b* = +90°, relative to straight ahead). Participants were instructed on the nature of the changepoint process, and they had the opportunity to learn the values of the task’s parameters (*H*_*cp*_, σ_*exp*_) in a practice session (not analysed). See section 4.1 for more details on the experimental procedure or consult the original publication by Krishnamurthy, Nassar, et al. (2017).

**Figure 2.**
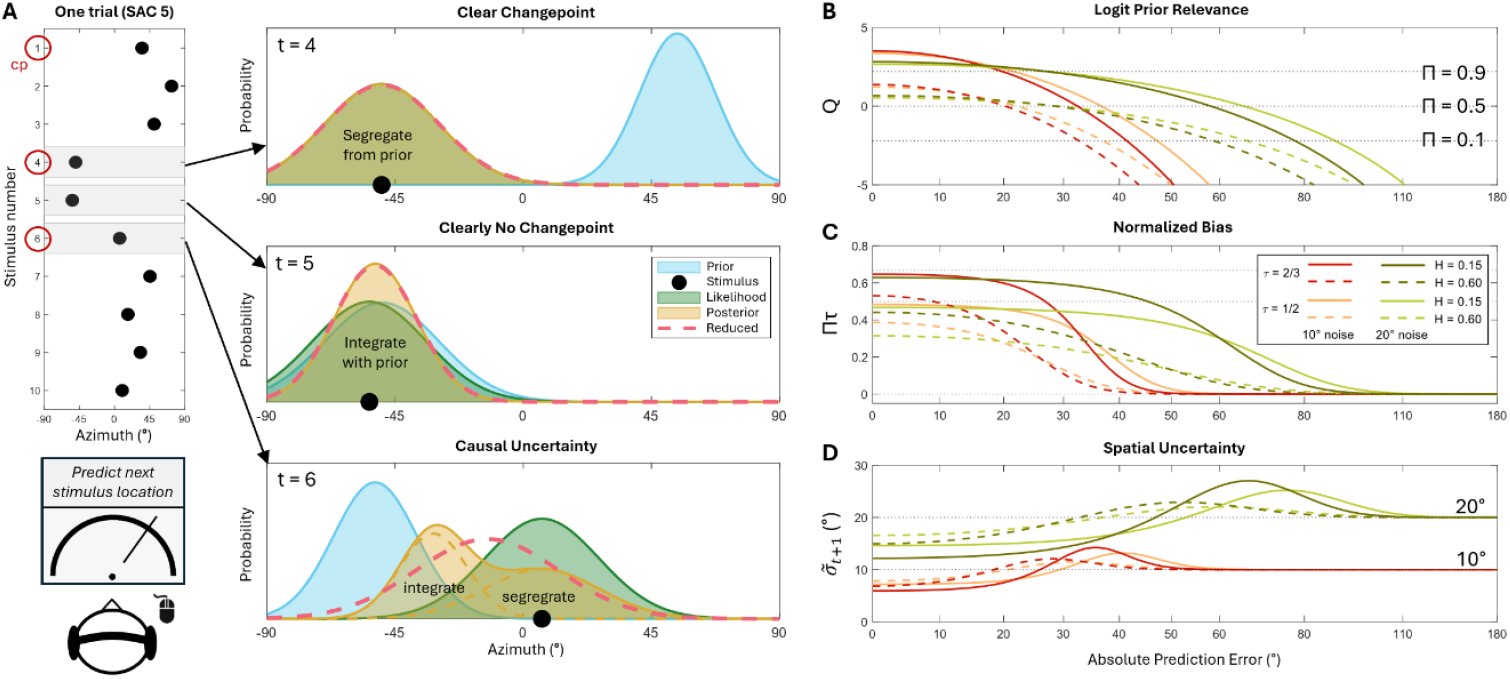
**The reduced Bayesian observer model puts the causality question centre stage after every stimulus: Did the latest observation originate from the same generative mean location as the prior (with added noise), or has there been a changepoint (cp)?** **A). Example trial in which an observer is presented with ten audiovisual stimuli. The generative mean changes twice after the start of the sequence: at *t*=4 and at *t*=6. Five consecutive stimuli are presented after the last cp (SAC 5), after which the participant responds with a prediction for the location of the upcoming stimulus. Ideally, the response location approximates the mean of the stimuli since the last cp. The sequential inference process to estimate this mean location is illustrated for three stimuli. Upon experiencing a large precision-weighted prediction error, a cp is inferred, and the prior becomes irrelevant for the current generative mean. So, the posterior (and next prior) is based on the likelihood only (*t*=4). Instead, small prediction errors indicate a low probability of a cp, thus likelihood and prior are integrated to improve precision of the mean estimate (*t*=5). But what to do when there is (causal) uncertainty about the occurrence of a cp (*t*=6)? The moderately sized prediction error suggests a cp, but the overlap between prior and likelihood indicates that they could have also originated from a common generative mean, i.e., *μ*_6_ = *μ*_5_. A fully Bayesian observer computes a posterior as a weighted mixture of two posterior components (dashed lines), each conditional on a causal hypothesis (cp or not). Alternatively, the reduced Bayesian observer summarizes that mixture distribution by its mean and variance, and thus simplifies the posterior (and next prior) to a single normal distribution.** **B-D). The prior relevance measure (*Π*) is key to the inference process of the reduced Bayesian observer. Panel B depicts its logit transformation (*Q*, i.e., the posterior log-odds of no changepoint) as a function of absolute prediction error 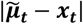, for two experimental noise conditions, two levels of prior reliability (*τ*), and two changepoint hazard rates (*H*_*cp*_). Panel C shows the resulting normalized bias towards the prior (at 1, relative to the last stimulus), which equals the product of prior relevance and prior reliability. Panel D illustrates how spatial uncertainty about the generative mean depends on the latest prediction error: it is low for small errors due to integration of prior and likelihood, it is identical to the experimental noise for large prediction errors, and highest in-between because of causal uncertainty.**

### 2.1 Reduced Bayesian observer model

Bayesian inference provides a statistically optimal method of dealing with the uncertainty that is caused by the combined presence of noise and changepoints in order to minimize the squared prediction errors. In section 4.3, we have derived the fully Bayesian solution for this task design (Adams & MacKay, 2007). However, since that ideal observer model relies on an infinitely large memory capacity and is computationally complex, we instead consider the reduced Bayesian observer model that was introduced by Nassar et al. (2012). This model presents a more plausible inference algorithm to be utilized by the brain and it has the additional benefit that it makes the causal inference process (i.e., changepoint or not) very explicit and insightful due to its simplicity.

Since the upcoming stimulus location *x*_*t*+1_ is equal to its generative mean *μ*_*t*+1_ plus some random noise, the reduced Bayesian observer makes predictions about *x*_*t*+1_ based on its best prediction for *μ*_*t*+1_: i.e., 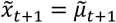. Here, the tilde indicates that these are predictions. To compute its prediction about the upcoming generative mean, the modelled agent attempts to compute the mean of the preceding stimuli since the last changepoint. If a changepoint occurred at the latest timepoint, *cp*_*t*_ = 1, then only the last stimulus is relevant, and the associated uncertainty is given by the learned estimate (indicated by circumflex) of the experimental noise:

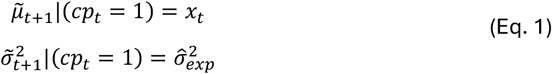

On the other hand, if there was no changepoint at timepoint *t*, then there are multiple relevant stimuli over which the agent should compute the mean. Fortunately, one does not have to remember all stimuli locations since the last changepoint. Instead, Bayesian observers compute the mean iteratively via reliability-weighted integration (which implies that they only have to remember the current mean and associated variance):

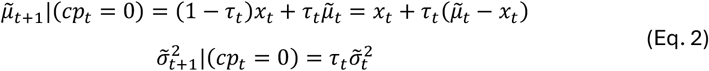

where the weight *τ*_*t*_, termed ‘prior reliability’, is computed as a relative measure of prior precision (i.e., reciprocal of variance):

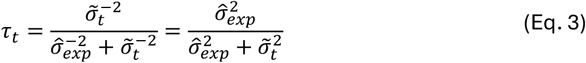

Notice how we expressed the predicted generative mean as a sum of the last stimulus location and a bias towards the prior. The normalized bias size (between 0 and 1, for predictions at *x*_*t*_ and 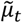, respectively) is equal to the prior reliability *τ*_*t*_. Notice further how the spatial uncertainty about the generative mean decreases with integration because 0 ≤ *τ*_*t*_ < 1. In the absence of changepoints, the spatial uncertainty would be reduced by a factor that is equal to the inverse of the number of observations (cf. standard error of the mean).

New evidence should be integrated with prior beliefs (Eq. 2) when the two relate to the same generative mean, and they should be segregated (Eq. 1) when they belong to different generative means, i.e., after a changepoint (Figure 2A). But how does one know whether a changepoint has occurred? A Bayesian observer infers this from the posterior probability of no changepoint, *Π*_*t*_, termed ‘prior relevance’ (Krishnamurthy, Nassar, et al., 2017). Since it is a posterior measure, *Π*_*t*_ depends on a combination of pre-existing beliefs and an evaluation of the newest observation. To be able to separate these two contributions, we present the formula for the posterior log odds of no changepoint, *Q*_*t*_, i.e., the logit transform of *Π*_*t*_:

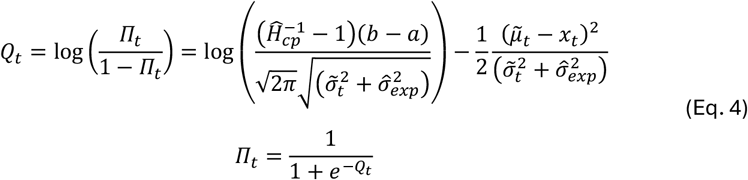

where the unbounded quantity *Q*_*t*_ is back-transformed to the posterior probability, 0 ≤ *Π*_*t*_ ≤ 1, via the monotonic logistic function.

The first term (within the logarithm) represents an a-priori belief about the prior’s relevance that is independent of the last stimulus location *x*_*t*_. The prior’s relevance decreases with a larger changepoint hazard rate estimate, 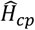, it increases with a larger spatial range of the generative mean (*b* − *a*), and it decreases with more uncertainty about the location of the upcoming stimulus 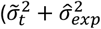, i.e., summed variance of prior on *μ*_t_ and experimental noise). The second term specifies by how much the (logit-transformed) prior relevance decreases as a function of the squared prediction error, i.e., after having observed *x*_*t*_ (Figure 2B). Importantly, a particular squared prediction error reduces the prior relevance by a smaller amount when the a-priori uncertainty about the stimulus location was larger. This precision-weighted squared prediction error term can thus be intuitively interpreted as a measure of surprise about the latest stimulus location *x*_*t*_ that is conditional on the prior’s relative location and uncertainty (see section 4.4).

Finally, the reduced Bayesian observer computes its prediction for the generative mean as a weighted combination of the two causal structures: changepoint or not. The weights are determined by the prior relevance:

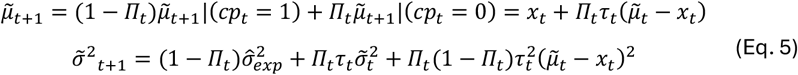

where we have once again expressed the prediction for the generative mean as a sum of the last stimulus location and a bias towards the prior. The normalized bias size is equal to the product of prior reliability *τ*_*t*_ and prior relevance *Π*_*t*_ (Figure 2C; see also section 4.5). Note that the predicted spatial uncertainty is formed by a weighted average of the variances of the two causal structures plus a third term that indicates the additional variance due to the causal uncertainty, *Π*_*t*_ (1 − *Π*_*t*_), which further depends on the squared prediction error and prior reliability. The contribution of this causal uncertainty term can be considerable. Because of it, the spatial uncertainty about the generative mean based on multiple stimuli can even exceed the spatial uncertainty based on a single stimulus 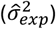 in situations where there is a high degree of causal uncertainty about whether or not a changepoint occurred (Figure 2D).

While it may seem counterintuitive to update one’s beliefs with the (weighted) average of two distinct possibilities (changepoint or not), thus placing one’s best prediction for upcoming stimuli somewhere in the middle where neither causal explanation matches particularly well, we emphasize that such concerns are unjustified. The prior relevance drops to near-zero for large prediction errors, so that the weighted average does not significantly bias one’s belief towards the prior (Figure 2C). Instead, the resulting prediction would be near the latest sensory evidence, in accordance with what one should expect after a noticeable changepoint. As the prediction error decreases in size, the prior relevance and resulting bias increase, and the updated belief will be exactly in the middle when *Π*_*t*_ *τ*_*t*_ = 0.5. However, as the prediction error decreases, we also effectively increase the amount of overlap between the two posterior component distributions, conditional on a changepoint or not (Eqs. 1-2; Figure 2A, dashed yellow lines in bottom panel). This means that the weighted average prediction 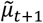 in such cases will actually be relatively well supported by both causal explanations.

### 2.2 Qualitative evaluation

In this section, we compare general patterns and trends in the predictions of the reduced Bayesian observer model (Eqs. 3-5) against the behavioral responses of the participants. To gain further insights about the model and participants’ perceptual decision making, we also contrast them against predictions of two other hypothetical observers.

The first hypothetical observer is naïve to the generative process of the stimuli sequences or chooses to ignore it. The naïve observer bases its prediction responses on the last observed stimulus location exclusively: 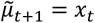. Within the computational framework of the reduced Bayesian observer this behavior can be achieved by setting the estimate of the changepoint hazard rate equal to one, 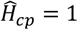, which results in a constant prior relevance of zero, *Π*_*t*_ = 0. The naïve observer is non-Bayesian because it never integrates new evidence with prior beliefs. Hence, it is unable to decrease its prediction errors after observing multiple stimuli from the same generative mean. However, one could argue that the naïve observer opts for a cost-effective strategy in an uncertain environment: it avoids complex computations entirely in exchange for moderately larger but unbiased prediction errors (Eissa et al., 2022).

The second hypothetical observer is omniscient with regards to the changepoints. In other words, this unrealistic observer has full knowledge over when changepoints occur and thus experiences no causal uncertainty. It correctly sets the prior relevance *Π*_*t*_ to either zero (*cp*_*t*_ = 1) or to one (*cp*_*t*_ = 0) and it subsequently updates its beliefs about the generative mean according to Eqs. 3 and 5. Predictions of the omniscient observer are thus based on the mean location of all observed stimuli since the last changepoint. The omniscient observer sets a comparative benchmark for performance under conditions without causal uncertainty.

In a first analysis, we compare the predicted generative means of the omniscient observer against participants’ prediction responses (Figure 3A). In general, they are highly correlated (Pearson’s *ρ* = 0.98 [0.96 – 0.99] and *ρ* = 0.95 [0.94 – 0.96], for the low and high noise conditions, respectively. N.b. we consistently report group-level median [Q1 – Q3]). There was only a very small bias towards the centre of space for the participants’ responses overall (regression slope = 0.96 [0.92 – 0.98] and 0.94 [0.90 – 0.98]). These minor biases are in accordance with the reduced Bayesian observer model, and they are predominantly caused by a regression to the mean due to partial integration with stimuli prior to unnoticed changepoints. Importantly, a fully Bayesian ideal observer model portrays much larger central biases, because it attempts to mitigate potentially large prediction errors in case of a changepoint at the upcoming timepoint by multiplying its estimate of the current generative mean by a factor that depends on the changepoint hazard rate (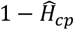, i.e., regression slopes < 0.85; see section 4.6). We conclude that participants did not adjust their predictions for the possibility of an upcoming changepoint, and instead they seem to have provided responses based on their belief about the current generative mean. This has been reported previously (Nassar et al., 2010) and was thus implemented in the reduced Bayesian observer model (and all other model variations that were tested here, see section 2.4).

**Figure 3.**
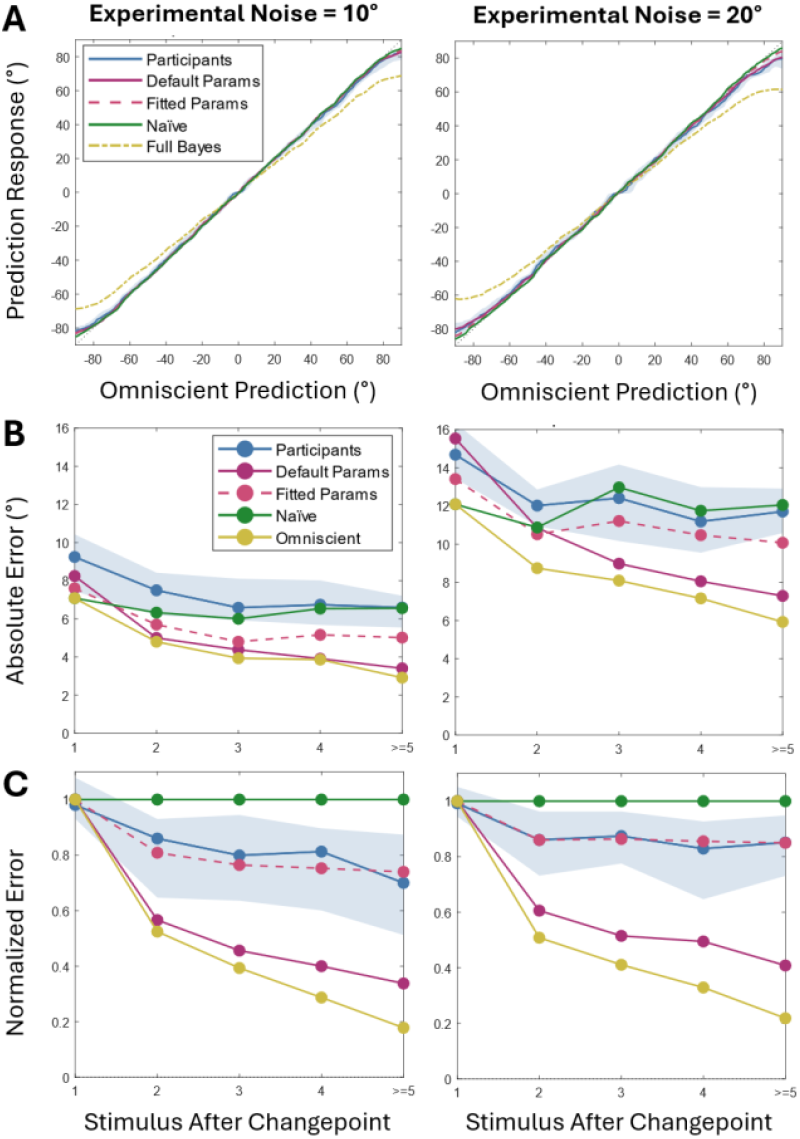
**Participants’ prediction responses are reasonably accurate, but the reduced Bayesian observer model with default parameters performs better.** **A). On average, participants’ responses correlate well with the omniscient observer model for the two experimental noise conditions (left and right panels). The fully Bayesian observer biases its predictions towards the centre of space to accommodate potentially upcoming changepoints. This bias is not present for participants, and it is therefore not modelled for the other observers (naïve, omniscient, and reduced Bayesian, with default parameters or with parameters that were fit to the participants’ data; see section 2.3).** **B-C). The response error relative to the true generative mean (unknown) decreases with larger SAC levels due to integration of consecutive stimuli, but participants’ absolute error remains larger than the naïve observer because of additional response noise (panel B). When the errors are normalized with respect to the naïve observer’s responses (at 1), then the random response noise averages out and the relative accuracy improvement becomes visible (panel C).** **The traces in all panels depict the group-level median of the individuals’ median response, per SAC bin (in B-C), or using a rolling kernel method (in A). Note that the rolling median results in small edge artifacts (panel A), with an apparent central bias for peripheral locations even for the naïve observer (and in addition to the integration-based bias for the other observers). The blue shaded region depicts the range between the group-level 25% (Q1) and 75% (Q3) participants.**

Perhaps the most important prediction that any Bayesian observer model makes is that its accuracy should (generally) improve by integrating the newest evidence with prior beliefs. We would thus expect the average absolute error to decrease in size as more stimuli are integrated. Figure 3B shows that this is certainly true for the omniscient observer, whose errors relative to the true generative mean *μ*_*t*_ decrease as the number of stimuli after [the last] changepoint (SAC) increases. Participants also reduced their errors relative to SAC level 1, but they did not improve further when more than two stimuli were observed after the last changepoint: A 2 × 5 repeated measures ANOVA shows a main effect of low vs. high noise, *F*(1,28) = 397, *p* < .001, and a main effect of SAC level, *F*(4,112) = 25.2, *p* < .001; post hoc tests with Holm correction confirm the differences between SAC 1 and all other levels, *p* < .001, but no other pairwise differences reach significance, *p* > 0.05.

So, participants reduced their errors when more than one stimulus was available after a changepoint. Nevertheless, they did not outperform the naïve observer that simply predicts the next stimulus at the location of the last stimulus. This may suggest that it would have been better for participants to adopt the same naïve strategy instead of attempting to infer when changepoints occur and integrate relevant stimuli. However, we argue that participants’ worse performance can be explained by additional random noise in the response phase, i.e., following the perceptual decision (see section 4.8, but not included in the models for this analysis). Such response noise could, for example, arise due to imprecision of the sensorimotor system or because of a self-imposed speed-accuracy trade-off. Support for the hypothesis of additive post-decision response noise comes from the analysis of errors that were normalized with respect to the naïve observer (Figure 3C). Single-trial normalization was achieved by dividing the actual error (*response* − *μ*_*t*_) by the difference between last stimulus and generative mean (*x*_*t*_ − *μ*_*t*_), such that a response at *x*_*t*_ receives a normalized error of 1 (constant for the naïve observer), while a response at *μ*_*t*_ naturally results in a normalized error of 0 (see section 4.11). Contrary to the absolute errors (Figure 3B), this procedure enables us to average out random response noise in the normalized errors. The analysis clearly demonstrates that participants reduced their average normalized errors relative to the naïve observer as they integrated evidence over multiple stimuli (sign tests: *p* < .005 for all SAC > 1 in both noise conditions, Bonferroni corrected).

The reduced Bayesian observer model predicts smaller errors (both absolute and normalized) than those of the participants. However, its performance is worse than the omniscient observer, thus illustrating the effect of causal uncertainty, i.e., the ambiguity about the veracity of changepoints. Errors will be larger if a true changepoint is misinterpreted as an outlier due to random noise (SAC 1), and vice versa, errors will also be larger if a changepoint cannot be ruled out after observing an actual outlier (SAC > 1). One can see that the reduced Bayesian observer struggles more with causal uncertainty in the high noise condition, as the difference with the omniscient observer’s performance is larger in the high noise condition as compared to the low noise condition, especially for low SAC levels. Note that the Bayesian observer performs worse than the naïve observer at SAC 1 (higher absolute error, Figure 3B), thus indicating a trade-off in adopting Bayesian causal inference: it improves performance overall, but it exacerbates errors after environmental changes. Participants, on the other hand, may have used a more conservative, risk-averse strategy in which they only moderately benefit from integration, but also attempt to avoid making large errors after changepoints.

Accuracy improvement with increasing SAC levels is achieved in the reduced Bayesian observer model by biasing its beliefs towards the prior, relative to the latest evidence. In Figure 4A we show that participants also exhibit such biases. Since it is impossible to know the covert prior beliefs of participants, we instead used the predicted prior beliefs of the omniscient observer (i.e., the mean of all stimuli since the last changepoint) to compute the normalized biases (for both participants and modelled data). This prior location is probably not precise, but it allows us to approximate the experienced prediction error for the latest observation within a reasonable margin, i.e., 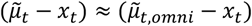. We thus computed participants’ normalized bias towards the prior as 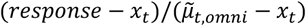, where a response at 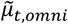 results in 1, and a response at *x*_*t*_ indicates no bias. Note that this normalized bias corresponds to the product of prior relevance and reliability, *Π*_*t*_ *τ*_*t*_, in Eq. 5 (although for the figure we computed the modelled biases in the same way as for the participants’ responses).

**Figure 4.**
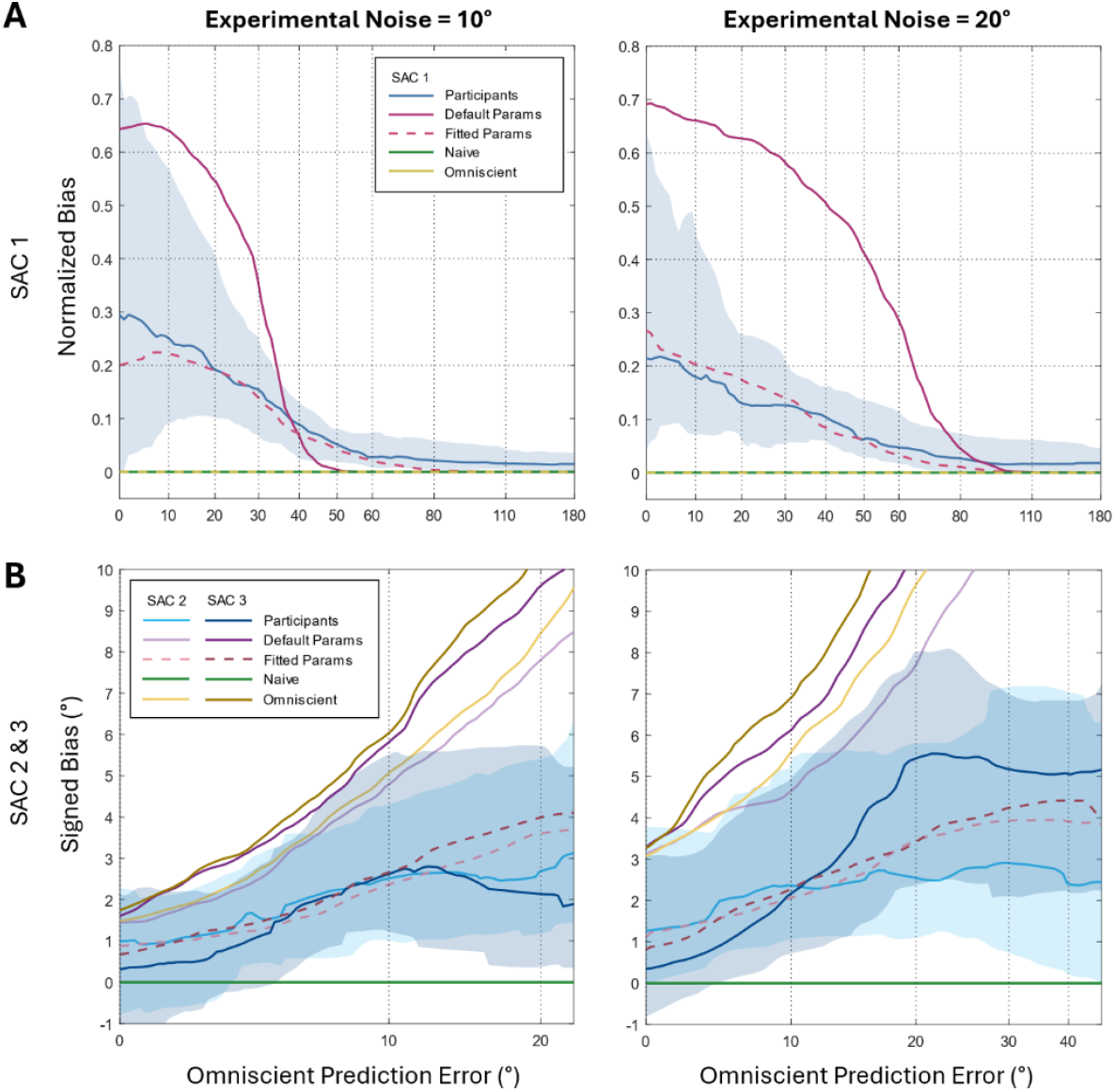
**Normalized bias towards the prior (at 1) for SAC 1 trials (panel A) and signed bias towards the prior (positive sign) for SAC 2 and SAC 3 trials (panel B), as a function of the experienced prediction error. The prior’s location was approximated by means of the omniscient observer.** **The traces depict the group-level median of the individuals’ local median response, as computed via a rolling kernel method. For appropriateness of that procedure, the prediction errors were non-linearly scaled to approximate a constant density of trials over the x-axis. Note that the rolling median results in small edge artifacts which cause the signed bias to appear larger than zero for the smallest prediction errors.** **The blue shaded region depicts the range between the group-level 25% (Q1) and 75% (Q3) participants. For the modelled observers we only depict the group-level medians (naïve, omniscient, and reduced Bayesian, with default parameters or with parameters that were fit to the participants’ data; see section 2.3).**

In Figure 4A, we have limited the analysis to SAC 1 trials only in order to clearly demonstrate the effect of increasingly large prediction errors. The pattern of a gradually decreasing normalized bias is similar to what is predicted by the reduced Bayesian observer model. This suggests that participants do indeed compute a prior relevance measure (Eq. 4) and use it to adaptively scale their bias towards the prior (Eq. 5). In doing so, they effectively computed a weighted average of the two causal structures (changepoint or not) and thus incorporated causal uncertainty into their updated belief. Furthermore, we also observe an effect of the experimental noise condition. Relative to the low noise condition, the maximal bias for the high noise condition is smaller for small prediction errors (Wilcoxon signed rank test on biases at 20° prediction error: *z* = -1.98, *p* = 0.048, low noise = 0.17 [0.10 – 0.42], high noise = 0.13 [0.06 – 0.27]), but the bias remains higher for moderately large prediction errors (Wilcoxon signed rank test on biases at 40° prediction error: *z* = 2.02, *p* = 0.043, low noise = 0.09 [0.02 – 0.13], high noise = 0.10 [0.07 – 0.14]). Both of these effects were predicted by Eq. 4 of the reduced Bayesian observer model via the noise dependence in the denominators of the a-priori relevance (peak bias for small predictions errors) and surprise term (reduction of relevance with prediction error size).

Next, we looked at the biases for SAC levels 2 and 3 (Figure 4B). Since the effect of small misestimations of participants’ prior (i.e., 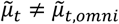) leads to excessive amounts of noise in the normalized biases at small prediction errors (also observed in the intersubject variance, see shaded area in Figure 4A), we instead focus here on non-normalized biases (*response* − *x*_*t*_). However, we modified the bias signs to ensure that a positive bias always points towards the predicted prior location 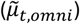. Importantly, we only included SAC 2 trials for this analysis if they were preceded by a large prediction error (>52.7°, as determined by a median split), such that the preceding changepoint had likely been noticed. Hence, the spatial prior for these SAC 2 trials should have been based on the preceding stimulus only. Likewise, we limited the analyses of SAC 3 trials to those trials that were associated with a large changepoint prediction error (>52.7°), followed by a relatively small disparity between the next stimuli (i.e., |*x*_*t*−2_ − *x*_*t*−1_ | had to be <13.7° or <27.4° for the low and high noise conditions, respectively; where the threshold values were based on excluding one third of the trials with the largest disparities). Hence, the spatial prior for these SAC 3 trials should have been based on the average of the two preceding stimuli, and the prior’s spatial uncertainty 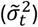 for these SAC 3 trials should thus be smaller than the prior’s uncertainty for the SAC 2 trials.

The general pattern of the signed biases in Figure 4B is similar as predicted by the reduced Bayesian observer model: the bias towards the prior increases in size with increasing prediction errors, but then starts to flatten and eventually decrease at larger prediction errors as a result of the smaller prior relevance (*Π*_*t*_). The peak of participants’ signed biases is approximately reached at a prediction error that is similar to the experimental noise, 10° and 20°. For larger prediction errors, participants apparently already start to question whether a changepoint has occurred. The reduced Bayesian observer model peaks later, at approximately 29° and 26° with low noise and 53° and 47° in the high noise condition, for SAC 2 and SAC 3 trials respectively. The predicted differences between the SAC levels occur because the spatial uncertainty of the prior 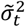 is smaller for SAC 3 than SAC 2, which increases the surprise (2^nd^ term in Eq. 4), thus lowering the prior relevance at smaller prediction errors. These predicted small differences are not clearly observed in the participants’ data. However, another model prediction is that the size of the biases are larger for SAC 3 than for SAC 2, independent of the prediction error, because of a difference in the prior reliability *τ*_*t*_ (Eq. 3). Participants show this effect in the high noise condition (Wilcoxon signed rank test on biases at 20° prediction error: *z* = 2.15, *p* = 0.031, SAC 2 = 2.31° [1.02° – 6.25°], SAC 3 = 5.40 [2.23 – 7.74]), but not in the low noise condition (Wilcoxon signed rank test on biases at 10° prediction error: *z* = 0.42, *p* = 0.67, SAC 2 = 2.67° [1.08° – 4.54°], SAC 3 = 2.34 [1.28 – 5.73]).

### 2.3 Parameter optimization

In the above analyses we have shown that hallmarks of Bayesian causal inference are present in the response patterns of the participants. However, quantitatively, the reduced Bayesian observer model with default parameters (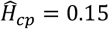 and 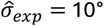 or 20°, for low and high noise conditions, respectively) does not show a very good fit. In particular, the model severely overestimates participants’ biases towards the prior. Assuming that the principal mechanisms for Bayesian causal inference are present in the brain (Sham & Beierholm, 2022), one plausible suggestion to reconcile the experimental results to the theoretical predictions is that participants were using incorrect estimates of the model’s parameters (Nassar et al., 2010). For example, larger changepoint hazard rate estimates, 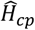, would have the effect of decreasing the prior relevance *Π*_*t*_ independent of the prediction error, thus resulting in smaller biases overall.

Participants were given the opportunity to learn about the changepoint hazard rate (*H*_*cp*_) and the amount of experimental noise (σ_*exp*_) in both conditions in a practice session. However, learning these parameters by merely observing the stimuli locations is no easy task (Wilson et al., 2010; Piray & Daw, 2021). A large prediction error may indicate that a changepoint occurred. If many changepoints are inferred, then the subjective hazard rate estimate should be larger. But prediction errors can alternatively be interpreted as due to random noise, in which case the subjective experimental noise estimate should be larger. Inference about the parameters of the generative process thus essentially revolves around a volatility (changepoint) vs. stochasticity (noise) trade-off, similar to the causal inference problem itself. Therefore, we would expect the two parameter estimates, 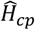 and 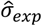, to be anticorrelated.

One method that the brain may apply to learn such ‘environmental parameters’ is based on an objective to minimize overall surprise (Friston, 2010). Without specifying how such a learning algorithm may work, we here computed the overall surprise that a reduced Bayesian observer would experience in the current experiment (i.e., summed over all stimuli) for a large number of predetermined parameter combinations (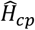 and 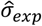) on a regular grid (see section 4.10). The predicted amounts of surprise are visualized as a background colour coding in Figure 5A. One can see that an agent with parameter estimates near the true values (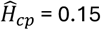 and 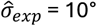 or 20°) minimizes the overall surprise. Some parameter combinations are predicted to lead to excessively high amounts of surprise (e.g., 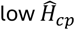 and 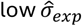), thus indicating that such parameter estimates would lead to a significant mismatch between the stimuli locations that are predicted and those that are experienced. However, for many parameter combinations near the true values, surprisal remains relatively low. This suggests that all such estimates are reasonably likely for near-Bayesian observers that effectively learn the statistics of their sensory environment. The expected anticorrelation between the hazard rate and experimental noise estimates is highlighted as a dashed line (indicating the value of 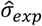 that minimizes surprisal for a particular value of 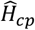).

**Figure 5.**
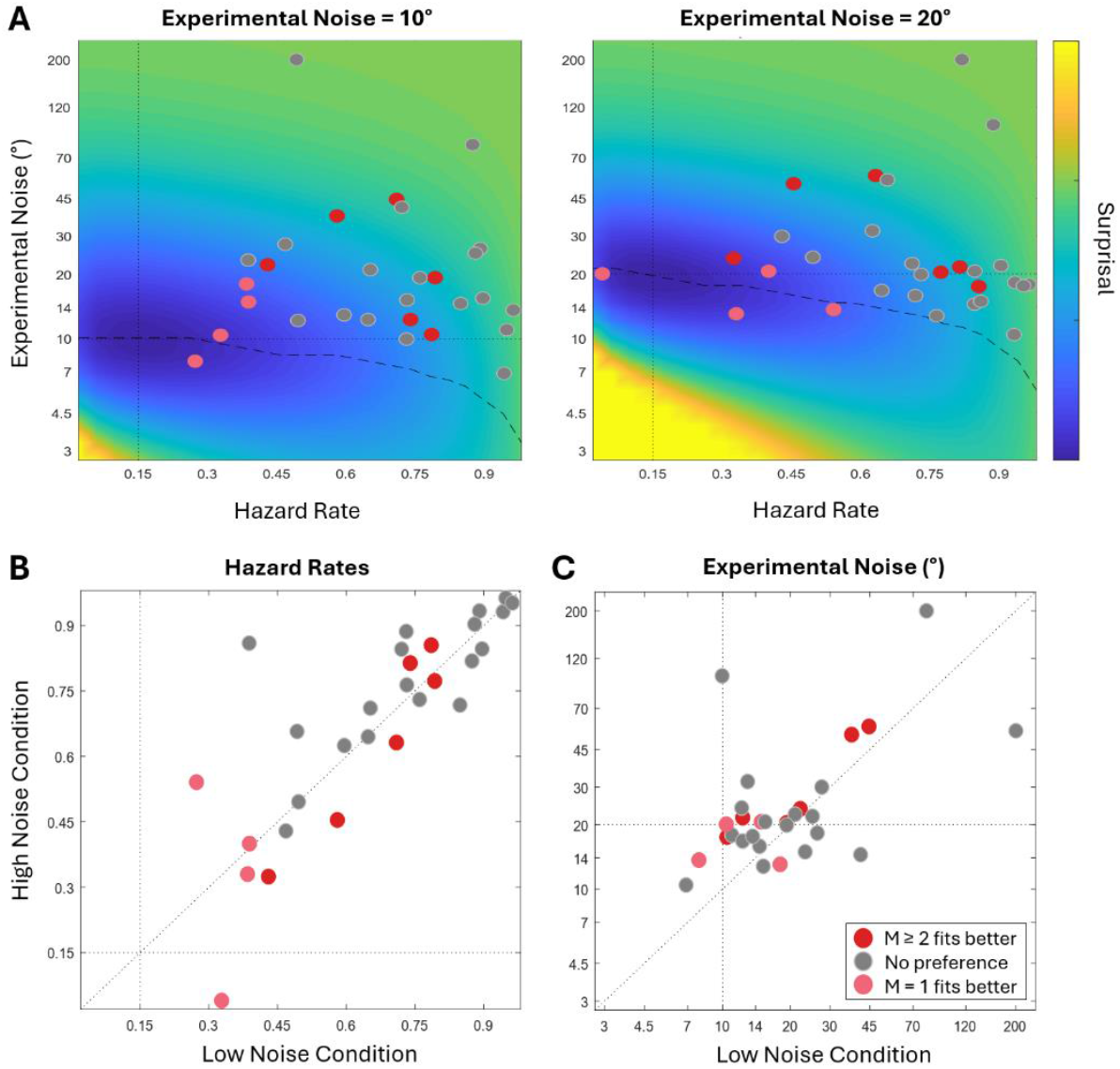
**Fitted parameter values of the reduced Bayesian observer model.** **Panel A. depicts the fitted changepoint hazard rate 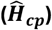 and experimental noise ^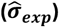^ estimates of the individual participants (separately for both experimental noise conditions, in left and right panel) on top of a modelled surface of overall (summed) surprisal levels that a reduced Bayesian observer would experience with such parameter values. Surprisal is minimized for parameter estimates that are equal to the true generative parameter values (thin dotted black lines). The dashed black lines illustrate the expected negative correlation between hazard rate and experimental noise estimates.** **Panels B and C show the same fitted hazard rates and experimental noise estimates, but now as a comparison of the noise conditions.** **Individuals’ colour coding depends on the model comparison results (section 2.4 and Figure 7B): dark red for participants that are better fit by models with a larger memory capacity, light red for participants that are better fit by the reduced Bayesian observer model with limited memory capacity, and grey otherwise.**

To obtain estimates of the parameters that participants may have used during the task we fitted the reduced Bayesian observer model to each set of responses from one individual and one experimental noise condition (by means of maximum likelihood estimation, see section 4.9). As expected by the relatively low biases (see Figure 4), we found that the fitted hazard rates for most participants were much higher than the true hazard rate (Figure 5B). While there was considerable variance of the fitted hazard rates across participants, the values per participant were quite consistent across the two experimental conditions (Pearson’s correlation coefficient ρ = 0.81). Furthermore, we found that the fitted experimental noise parameters (Figure 5C) were reasonably accurate for most participants, though not for all, and they were smaller in the low noise condition than in the high noise condition (low noise 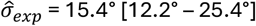, high noise 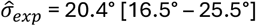, Wilcoxon signed rank test: *z* = -2.17, *p* = 0.030). Interestingly, we did not observe the predicted negative correlation between the fitted hazard rates and log-transformed experimental noise parameters (Pearson’s *ρ* = -0.04 and *ρ* = -0.02, for low and high noise condition respectively, but these correlations were not significant: *p* > 0.8). In fact, most of the largest experimental noise parameters were for participants whose hazard rate estimates were also large (Figure 5A). Those participants showed overall small biases (large 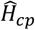) that were nonetheless non-zero even at large prediction errors (large 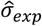).

Allowing the changepoint hazard rate and experimental noise parameters to be fitted freely significantly improved the quality of the fits in terms of the coefficient of determination: R^2^ = 0.97 [0.92 – 0.98] and R^2^ = 0.94 [0.88 – 0.97], for low and high noise conditions, respectively. This is marginally higher (∼1% point) than the reduced Bayesian observer model with default parameter values, i.e., with 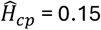 and 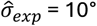 or 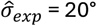 (Wilcoxon signed rank tests: *z* > 4, p < .001, for both conditions), and equally better than the naïve observer model (*z* > 4, *p* < .001, for both conditions). A more informative measure for the model’s improvement in quality of fit is the amount of explained variation in participants’ errors (*response* − *μ*_*t*_, i.e., as in Figures 3B and 3C). The group-level medians are R^2^ = 0.45 vs. 0.36 vs. 0.28, and R^2^ = 0.67 vs. 0.53 vs. 0.57, for the three models (reduced Bayesian model with fitted parameters, vs. with default parameters, vs. the naïve observer model) in the low and high noise conditions, respectively. N.b. remaining response variance was explained by the fitted reduced Bayesian observer model by means of occasional lapses (lapse rates *λ* = 0.017 [0.008 - 0.045] and *λ* = 0.043 [0.021 - 0.063], for low and high noise conditions, respectively) and by random response noise (σ_*resp*_ = 5.64° [3.65° – 8.26°] and σ_*resp*_ = 5.93° [3.90° – 11.0°]). Nevertheless, depicted predictions for the model with optimized parameter values in Figures 3 and 4 (dashed lines) do not include such lapses or response noise.

### 2.4 Model comparison

The reduced Bayesian observer model is a simplification of the fully Bayesian observer model. While the latter is unrealistic because of its complexity, the former is certainly not the only plausible inference algorithm. Here, we will compare the reduced Bayesian observer model against variations of itself that rely on different modelling assumptions.

The most striking simplification of the reduced Bayesian observer model is its limited memory requirement. The prior distribution at any timepoint is described by merely two parameters: the mean and uncertainty estimates about the generative mean (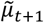 and 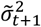, respectively). That prior distribution approximates the weighted mixture of the two conditional prior distributions for each causal structure with regards to the preceding stimulus: changepoint or not. The fully Bayesian observer model (Adams & MacKay, 2007) does not compute such approximate summary distributions. Instead, it remembers the conditional distributions and their associated weights (i.e., relevance). The advantage is that the weights of the conditional distributions can still be adjusted after new observations become available. For example, an agent may be uncertain about a possible changepoint at first, but this uncertainty may disappear if the next observation returns to a location that fits better with the previous prior. In that case, the spatial prediction of a fully Bayesian observer would correctly be dominated by the mean of all relevant stimuli, despite the temporary causal uncertainty that accompanied the penultimate stimulus. On the other hand, the memory-constrained reduced Bayesian observer’s prediction would generally be (slightly) less accurate (and more uncertain) because the observations are not weighted optimally: weight cannot be returned to observations whose relevance was once questioned (Figure 6A).

**Figure 6.**
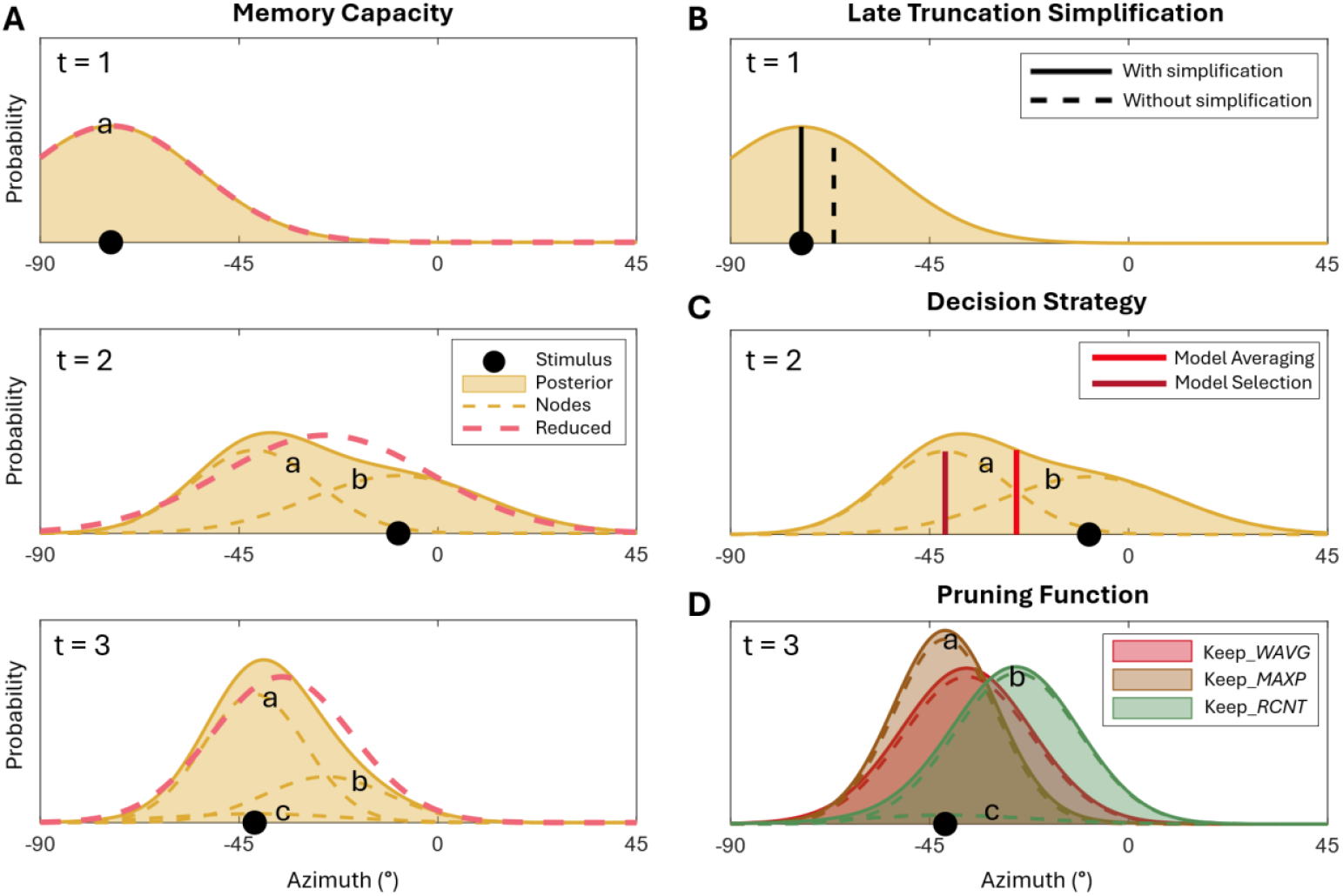
**Illustration of the modelling framework with four factors: memory capacity (A), late truncation simplification (B), decision strategy (C), and pruning function (D).** **A). The three panels depict a sequence of three stimuli (top to bottom) that illustrate the inference difference between a reduced Bayesian observer (M = 1, posterior depicted as dashed red line) and a similar Bayesian model with larger memory capacity (M ≥ 3, solid orange line with shaded area). Posterior distributions of models with extended memory consist of a weighted mixture of multiple nodes (dashed orange lines with a, b, and c letter indicators). The node’s weight indicates its relative posterior relevance for the inferred location of the generative mean. In this sequence, the second stimulus leads to high causal uncertainty (nearly equal weight for both nodes: a. latest cp at *t=1*, b. latest cp at *t*=2), but the third stimulus increases the inferred relevance of the node that codes for the hypothesis that no changepoint took place (a): i.e., it is most probable that all three stimuli stem from the same generative mean, and it is least probable that a changepoint took place at *t*=3 (c). The reduced Bayesian observer cannot retrospectively reassign weights to nodes and is therefore slightly less accurate (here: small bias towards penultimate stimulus) and less precise (larger spatial uncertainty) than a near-Bayesian observer with larger memory capacity.** **B). The late truncation simplification implies that the space boundaries are ignored until a response has to be given. At that point, the observer’s best prediction is moved to within the generative mean range, only if necessary. Without the simplification, observers compute the expectation (i.e., mean) of the truncated normal distribution as their best prediction, and this will bias their response towards the centre of space. Here, the difference is illustrated for a response at *t*=1 (from panel A). With the simplification, the observer responds to the mean of the (non-truncated) normal distribution: subsequent movement of the intended response location to within the generative mean range is unnecessary in this example.** **C). Whenever the posterior distribution consists of more than one node, the observer needs to decide on how to make a response (here depicted for *t*=2). The model averaging decision function computes the expectation of the weighted mixture distribution as its best prediction, thus essentially averaging two models of the world (i.e., nodes a and b). Instead, the model selection decision function deterministically selects the node with the highest weight (i.e., inferred relevance) as a base for its prediction response.** **D). The node pruning function determines how an observer satisfies the memory capacity limit *M*. Whenever the number of posterior nodes exceeds the memory capacity, the pruning function ensures that the prior for *t+1* consists of no more than *M* nodes. Here, this is depicted for the case at *t*=3, with a memory capacity of M=2. The keep_*WAVG* pruning function merges the oldest two nodes (a and b) by computing a weighted average node (while the newest node, c, for a cp at t=3, remains in memory as well, despite its small weight). The keep_*MAXP* pruning function keeps the node with the highest weight of the oldest two nodes and discards the other (i.e., node a is kept, b is discarded), whereas the keep_*RCNT* pruning function always keeps the most recent of the two oldest nodes (i.e., node b is kept, a is discarded). The node that is kept in memory also receives the weight of the discarded node, such that it now represents the hypothesis of a cp at t ≤ 2.**

So, being able to juggle multiple competing hypotheses simultaneously is beneficial for task performance. Moreover, it seems intuitive that humans are able to remember the approximate locations of at least a few recent stimuli, while also maintaining uncertainty about the presence of recent changepoints. Therefore, one of the primary factors of interest in our model comparison was the model’s memory capacity. We compared models whose predictive mixture distributions consisted of up to four conditional distributions, i.e., memory capacity *M* ∈ {1,2,3,4}. These conditional distributions are often referred to as ‘nodes’ of the full mixture distribution (e.g., Wilson et al., 2010, where the terminology is based on a graphical model depiction). Here, each node is represented by three parameters: its mean, variance, and weight (see section 4.3).

The reduced Bayesian observer model limits the memory capacity to *M* = 1 by summarizing its two conditional distributions via a weighted average (Eq. 5) after every observation. Essentially, it thus merges two nodes into one at every timepoint. We can flexibly extend the model’s memory capacity by merging progressively older nodes. The newest nodes, that are conditional on the hypothesis of a recent changepoint, and whose contributing stimuli are still fresh in memory, remain unchanged. For example, a model with a memory capacity of *M* = 4 would summarize its oldest two nodes such that the merged node is associated with the hypothesis that the last changepoint occurred four *or more* stimuli ago, while the latest three nodes are also kept in memory (see section 4.5).

Besides merging nodes via a *weighted average* (i.e., keep_*WAVG* node), one can also constrain the memory requirements via different rules (Figure 6D). For example, it has been suggested that humans may simply forget the oldest node once memory capacity has been exceeded (Skerritt-Davis & Elhilali, 2018). In that case an agent would only remember evidence that is associated with the hypotheses of a changepoint at any of the *recent* timepoints (keep*_RCNT*). A third option that was proposed previously (Adams & MacKay, 2007) is to discard nodes based on their inferred relevance (i.e., the nodes’ weights). In our implementation here, we keep either of the two oldest nodes, whichever has a *larger probability* of being relevant (keep_*MAXP*). We refer to these three procedures as node pruning functions (Wilson et al., 2010). Note that a model with the keep_*RCNT* pruning function and with minimal memory capacity, *M* = 1, is identical to the naïve observer model that we introduced in section 2.2.

Whenever the prior distribution consists of more than one node at the end of a trial (*M* > 1), the observer needs to decide on how to form a single prediction response out of the mixture distribution (see section 4.6). A Bayesian observer that attempts to minimize its squared errors computes a weighted average of the nodes’ means as its best prediction for the generative mean. Since each node is associated with a particular model of the world (i.e., causal structure), specifying when the last changepoint occurred, this decision strategy is known as ‘model averaging’ (Wozny et al., 2010). The consequence of model averaging is that an agent accepts to be somewhat wrong about the state of the world most of the time, but it avoids occasionally making excessively large errors. However, different strategies are also possible (Figure 6C). For example, under the ‘model selection’ decision strategy, an agent would instead attempt to maximize its accuracy most of the time, while occasionally accepting a large error. This agent selects the mean of the node with the largest weight (i.e., relevance) as its best prediction for the location of the generative mean.

A final factor that we consider in our model comparison is related to the computational complexity that arises as a result of the finite spatial interval of the generative mean *μ*_*t*_ (i.e., here between -90° and 90°). Bayesian observers account for the interval when they update the weights of the nodes by means of a complex computation that depends on the distance of the stimulus location to the interval’s boundaries (section 4.3) and again in the response decision phase when they compute the spatial expectations of the nodes (section 4.6). However, these complex computations can be omitted at the cost of a limited accuracy loss by only considering the interval constraint at the very end of the decision-making process. Under the late truncation simplification hypothesis (Figure 6B) spatial predictions are initially formed using simplified computations which do not account for the locations of the stimuli relative to the spatial boundaries, but the final prediction is moved to the nearest boundary if it would otherwise be outside of the generative mean interval (section 4.7). Note that the reduced Bayesian observer model makes use of the late truncation simplification.

In the above, we have described a framework of near-Bayesian models in which we combine the following factors: 1) memory capacity, 2) node pruning function, 3) decision strategy, and 4) late truncation simplification. In total, we fitted 42 model variations. There were 6 models with a minimal memory capacity (*M* = 1), one for each of the three pruning functions (keep_*WAVG*, keep_*MAXP*, and keep_*RCNT*) with and without the late truncation simplification. The other 36 model versions with larger memory capacity were organized factorially for *M* ∈ {2, 3, 4}, the three pruning functions, two decision strategies, and with and without truncation simplification. Since we fitted each model separately to the responses of both experimental noise conditions (to be able to compare the fitted parameters independently, see Figure 5), there were 84 model fits per subject. The models’ abilities to predict participants responses were compared by means of the log model evidence (*lme*; higher values indicate better fits), which is based on a sum of the maximized log likelihoods of the two conditions (see section 4.9).

Figure 7A depicts the group-level results for a fixed effects analysis (i.e., summed *lme* over participants). That analysis serves as a useful overview of the relative differences in predictive performance of all the models. However, we will draw conclusions from a selection of family-wise random effects analyses that are more robust to outlier subjects by allowing for group-level heterogeneity (Stephan et al., 2009; Rigoux et al., 2014; Acerbi et al., 2018). First, we focus on a comparison of the six models with *M* = 1. A 3 × 2 factorial analysis revealed a strong preference for the keep_*WAVG* pruning function (protected exceedance probability *pxp* = 1.00 for keep_*WAVG vs. 0*.*00* for both keep_*MAXP* and keep_*RCNT*) and a moderate preference for the late truncation simplification (*pxp* = 0.73 vs. 0.27). Next, we examined the 36 larger memory capacity models in a 3 × 3 × 2 × 2 factorial analysis. The result confirmed the preferences for the keep_*WAVG* pruning function and the late truncation simplification (*pxp* = 0.93 for keep_*WAVG* vs. 0.04 for keep_*MAXP* vs. 0.03 for keep_*RCNT*; and *pxp* = 0.96 vs. 0.04 in favour of the simplification). Moreover, the model averaging decision strategy seems to be crucial for larger memory models to fit participants’ responses well (*pxp* = 0.97 for model averaging vs. 0.03 for model selection). The fixed effects analysis, as depicted in Figure 7A, demonstrates that the choice of pruning function loses importance as the model’s memory capacity increases, while the model-averaging decision strategy becomes increasingly decisive as more nodes are available for a response decision.

**Figure 7.**
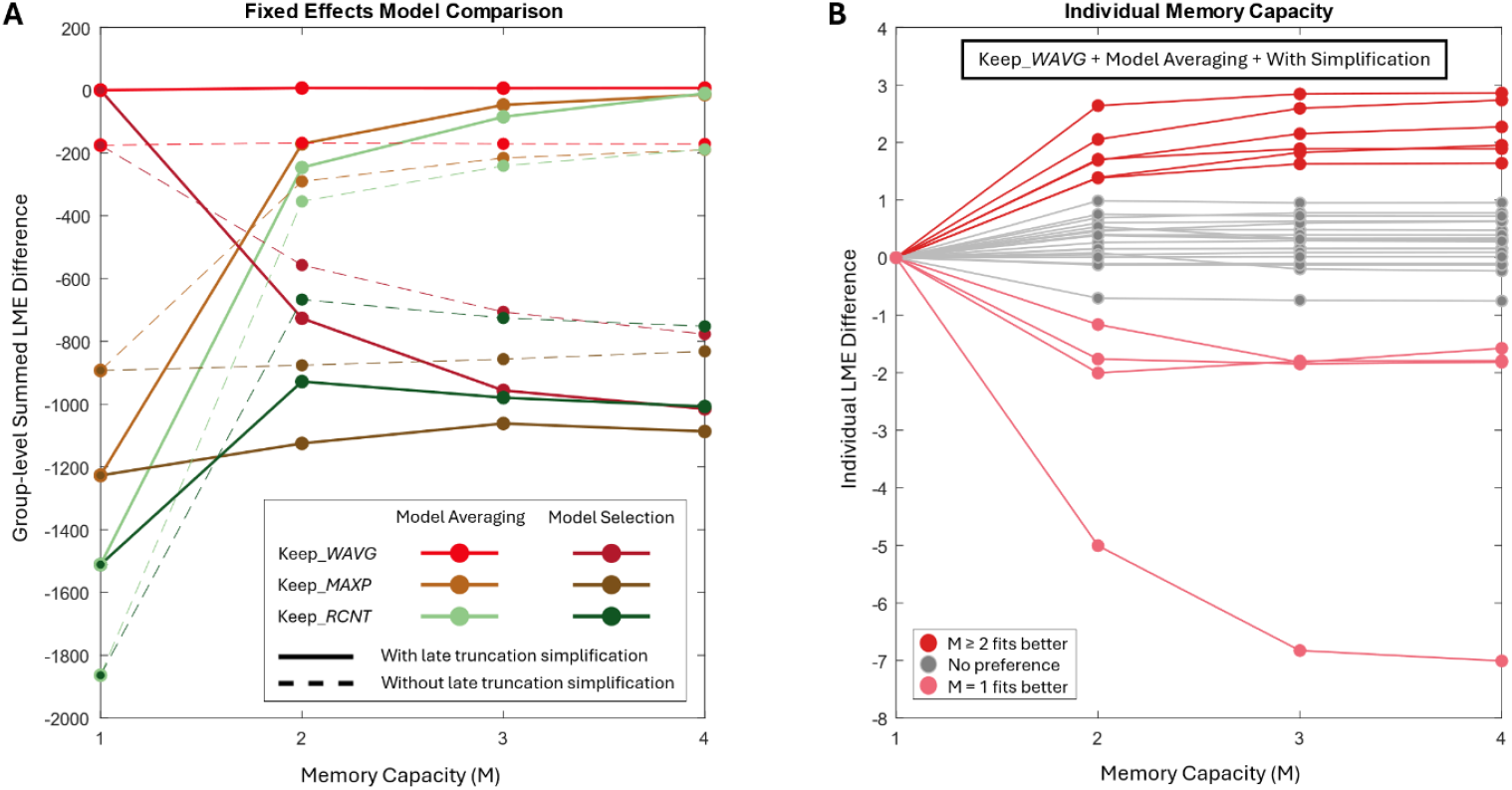
**Model Comparison Results. Panel A shows the fixed effects model comparison for the 42 models. The y-axis denotes the log model evidence (*lme*) of each model, summed across participants, relative to the reduced Bayesian observer model (memory capacity *M* = 1, keep_*WAVG* pruning function, model averaging decision strategy, and with late truncation simplification). The *lme* difference is dominated by the pruning function (colour coding) at low memory capacity (x-axis), while it is dominated by the decision strategy (light vs. dark) at larger memory capacities. Models with the model averaging decision strategy fit participants’ data better with the late truncation simplification, while models with the selection decision strategy generally fit better without the simplification. Panel B zooms in on the individual results of the memory capacity factor (x-axis), for models with keep_*WAVG* pruning function, model averaging decision strategy, and with late truncation simplification. Each line depicts the *lme* differences for one participant. The colour coding separates participants that were better fit by models with larger memory capacity (dark red), and participants who responded more like reduced Bayesian observers (light red), from participants without a clear preference (grey).**

To be able to fairly compare all four memory capacity settings against each other, we selected the four models that incorporated the late truncation simplification, the keep_*WAVG* pruning function, and the model averaging decision strategy (i.e., the preferred settings based on the previous analyses). The comparison was inconclusive, with an almost equal preference for each memory capacity (*pxp* = 0.25 vs. 0.24 vs. 0.25 vs. 0.26, for M from 1 to 4, respectively). The ambiguous result can be explained by a combination of two reasons that become clear when we plot the log model evidence of these four models for each individual (Figure 7B). First, most participants don’t seem to show a significant preference for any of the memory capacity settings (*lme* differences < 1, i.e., corresponding Bayes Factors < 3). Second, there is some heterogeneity in the population with six participants that are better fit by the larger memory capacity models, whereas the reduced Bayesian observer model (*M* = 1) is the best-fitting model for four other participants. Figure 7B also demonstrates that there are only minimal differences in the goodness-of-fit between the three larger memory capacity models. This is because these models make very similar predictions. So, while a memory extension to *M* = 2 is preferred to *M* = 1 by some participants (but not by others; a direct comparison is non-decisive: *pxp* = 0.31 vs. 0.69 in favour of *M* = 2), adding more model complexity through further memory extensions (*M* > 2) is not warranted by these results.

Finally, we note that there were only minimal differences between the fitted parameter values of the reduced Bayesian observer model (*M* = 1) and its model variation with larger memory (*M* = 2). The Pearson correlation coefficients, across participants, were *ρ* = 0.989 and *ρ* = 0.988 for (log-) experimental noise parameters, and *ρ* = 0.997 and *ρ* = 0.998 for the hazard rates (low and high noise conditions, respectively). Interestingly, we noticed a trend in the fitted parameter values of the four participants with a preference for the *M* = 1 memory setting: they all had relatively low hazard rates (<0.55, for both conditions; Figure 5B). This may potentially be indicative of individual differences in decision strategies. However, we are cautious about any such conclusion given the low number of participants with a clear model preference.

## 3. Discussion

In the current paper we have analysed the problem of changepoint detection in noisy environments from a perspective of Bayesian causal inference (Shams & Beierholm, 2022). The fundamental question that observers face is whether novel sensory signals should be integrated with prior beliefs to improve precision, or whether an environmental change has occurred that has rendered the prior irrelevant. The reduced Bayesian observer model (Nassar et al., 2012) puts this causality question centre stage after every new observation (Eq. 4) and then summarizes the probabilistic result to keep complexity at a minimum (Eq. 5). As such, it provides an elegantly simple algorithm for iterative Bayesian causal inference that can be utilized to estimate latent sources of consecutive sensory signals in volatile environments.

We tested the model’s performance in explaining human perceptual decisions on a behavioral dataset wherein participants made prediction responses after having been presented a sequence of noisy observations from static sources with occasional changepoints (Krishnamurthy, Nassar, et al., 2017). Our analyses brought forth three main findings: 1. The reduced Bayesian observer model predicted the response patterns qualitatively well. 2. It required freely fitting the model’s parameters to also quantitatively fit well. 3. Model comparison favoured the reduced Bayesian observer model over many model variations, but it was inconclusive with regards to the model’s memory capacity. In what follows, we shall discuss these three findings one by one and relate them to relevant literature in general and previous reports of learning in volatile environments in particular.

### 3.1. Hallmarks of Bayesian causal inference

We have shown that observers were partially able to average out noise from sequential observations during ongoing stimulus streams and thereby improved the accuracy of their predictions when more than one stimulus was available to base their decision on. In line with findings from cue integration experiments, they achieved this by weighing new sensory evidence versus existing beliefs according to their relative reliabilities (Eq. 3; prior attraction is larger for SAC 3 than for SAC 2 in the high noise condition). However, reliability is not the only factor that determines integration weights (Meijer et al., 2019; Rahnev & Denison, 2018). Since observers found themselves in a volatile environment, wherein prior information was not always relevant for their predictions, they dynamically reduced the prior’s weight in accordance with its inferred relevance.

According to the reduced Bayesian observer model, the weight of the prior is determined by the product of prior reliability and prior relevance (Eq. 5). The latter depends on an a-priori belief about the prior’s relevance as well as a surprise-based posterior evaluation (Eq. 4). In agreement with the model’s predictions, we found that reliance on the prior was larger in the low experimental noise condition than in the high experimental noise condition, at least for relatively small spatial disparities between the prior’s prediction and the latest observation. As the prediction error grew larger, the prior’s inferred relevance declined, and so we saw a gradual decrease of the normalized bias towards the prior (i.e., a measure of the prior’s weight). Moreover, the biases decreased faster in the low noise condition, because moderate prediction errors result in a larger surprise when the prior’s predictive precision is larger. Vice versa, the experimental condition with more stochastic noise led to flatter curves, thus less modulation of the prior’s weight as a function of the prediction error.

Smooth transitions from a large bias towards highly relevant priors, to near-zero biases for irrelevant priors, as we observed here, are modelled via a weighted average between two causal structures (i.e., models of the world that explain the causes of sensory signals; see Figure 1). In the reduced Bayesian observer model with limited memory (M=1) weighted averaging is established via the keep_*WAVG* pruning function, while for models with larger memory capacities this occurs primarily via the model averaging decision strategy. Regardless, models that compute a weighted average over the causal structures explained participants’ responses significantly better than models that did not (e.g., model selection decision strategy). By computing a weighted average, an observer essentially accounts for the uncertainty about the inferred causal structure. In the case of the reduced Bayesian observer model, the causal uncertainty is also literally incorporated into the updated belief about the source location via an additive term for the spatial uncertainty (Eq. 5; Figure 2D). This effectively ensures that appropriately low reliability is attributed to the updated prior at the next timestep if there was considerable uncertainty about the occurrence of a changepoint at the current timestep.

In the learning literature there has been a long-standing debate about the use of the prior relevance measure as a flexible moderator for the extent to which beliefs should be updated after a new observation. In that field it is common to speak of learning rates (taking perspective from the current belief) instead of biases towards the prior (from the perspective of the latest sensory signal), but in the current changepoint paradigm they are mathematically interchangeable: the momentary learning rate (*α*_*t*_) is equal to one minus the normalized bias (i.e., *α*_*t*_ = 1 − *Π*_*t*_ *τ*_*t*_). In the seminal delta rule model by Rescorla & Wagner (1972) learning rates **were assumed to be constant. However, later models, e.g**., **by Pearce & Hall (1980), allowed** learning rates to adjust flexibly to the size of the prediction error (Lee et al., 2020). As discussed, our results indicate that learning rates are indeed flexibly adjusted by the interaction of prior reliability and prior relevance (Figure 4).

Nevertheless, there are other recent studies that report a preference for a simpler heuristic with a constant learning rate (e.g., exponential decay or leaky accumulator models; Ryali et al., 2018; Norton et al., 2019). We believe that many such contradictory findings can be explained by the experimental task design which may not provide sufficient variety in the surprise term of the prior relevance, such that the learning rates only vary insignificantly even if an optimal adaptive strategy is applied (i.e., observations are information-poor; Ryali et al., 2018). For example, this may be the case in tasks that require observers to estimate a probability or to weigh two alternatives probabilistically. In such tasks, a single observation cannot be decisive evidence for a changepoint (or not), so the prior relevance does not take extreme values near zero (or one). This will also be the case for experiments with relatively large amounts of stochastic noise (e.g., experimentally induced, but potentially also due to sensory noise). As explained above, the prior relevance curve of an ideal observer in more noisy conditions will be relatively flat (as a function of prediction error). Under such high noise conditions, it may be difficult to conclude whether observers employ causal inference because the measured biases will appear very similar to a learner with a fixed learning rate (see also Heilbron & Meyniel, 2019). Moreover, we cannot exclude the possibility that humans choose to use a simpler cost-effective heuristic in a high uncertainty environment in which more complex solutions do not substantially improve accuracy (Tavoni et al., 2022; Verbeke & Verguts, 2024). Nonetheless, it is worth emphasizing that our results strongly suggest that changepoint designs with a moderately large range to noise ratio, such as the task that was used here (Krishnamurthy, Nassar, et al., 2017), introduce highly dynamic learning where observers adapt their use of the prior in accordance with its inferred relevance, thus confirming the key principle of Bayesian causal inference.

### 3.2. Reduced reliance on prior

We found that the reduced Bayesian observer model generally overestimated the biases towards the prior when its parameters were set to the correct values, i.e., the experimental noise and changepoint hazard rate that were used to generate the stimuli sequences. In other words, we reproduced previous reports that humans systematically used higher learning rates than an ideal observer would have done (Nassar et al., 2010; Lee et al., 2020, Bakst & McGuire, 2023). One may hypothesize that lower-than-ideal bias sizes can be explained by misestimation of the changepoint hazard rate. Specifically, if observers expect there to be more changepoints, then this would reduce the a-priori relevance of their priors. However, if participants did indeed overestimate the changepoint hazard rate, then we hypothesized that their experimental noise estimates should also be smaller than the true value. This is because of the expected trade-off between noise and changepoint inference rates for relatively large prediction errors (Payzan-LeNestour & Bossaerts, 2011; Piray & Daw, 2021).

We proceeded by fitting the reduced Bayesian observer model to the response data. The optimized hazard rate parameters were indeed (much) larger than the true changepoint rate, although there were considerable individual differences. Larger hazard rates reduced the modelled prior relevance, and therefore the biases, thus resulting in improved model fits to the response data. However, the optimized noise parameters were reasonably accurate; they were certainly not smaller than the true values, as would be expected from the presumed trade-off between experienced volatility and stochasticity. We interpret the lack of a trade-off as an indication that the fitted hazard rate parameters are not illustrative for the rate with which participants experienced changepoints. Instead, we believe that the optimized parameter values signify a reduced a-priori trust in the relevance of the prior, or a strategic choice to not rely on the prior as much as the reduced Bayesian observer. In other words, the probability 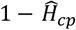 (or the non-negative quantity 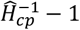 in Eq. 4) should rather be viewed as an individual’s tendency to bind novel sensory signals with existing beliefs. Similar binding tendencies, their plasticity and individual differences are an active area of research in the field of sensory cue integration (Quintero et al., 2022).

The lower-than-ideal binding tendencies for consecutive cues are in curious contradiction with the fact that human observers are often biased towards preceding stimuli and responses, also in experiments when there is no consistent relationship between successive stimuli or trials (thus when there is no benefit of such a bias). One explanation for these so-called serial dependence biases is that observers have learned that the world is generally stable, thus prior beliefs persist to stabilise perception even when objectively irrational within a rapidly changing experimental context (Kiyonaga et al., 2017). So, why would observers not make full use of prior information in experiments where it is actually useful to do so (like here)?

One plausible reason may be specific to the artificially volatile environment in changepoint paradigms. This may prompt participants to explicitly focus on the causality question (changepoint or not), thus making them overly attentive for deviations from stability. Relatedly, multisensory cue integration experiments have demonstrated that tasks that require explicit causal decisions result in more segregated percepts as compared to implicit causal inference tasks (Acerbi et al., 2018). Hence, being very aware of the constant potential for a changepoint could markedly reduce the extent to which one integrates consecutive sensory signals, independent of the actual hazard rate. By focusing on not missing any of the changepoints, participants may have adopted a suboptimal strategy for dealing with the changepoint versus noise trade-off. Bayesian ideal observers occasionally falsely infer ‘no changepoint’ (on SAC 1 trials), and so they bias their responses towards an irrelevant prior. In exchange, they more often than not correctly integrate successive stimuli (SAC > 1) and thus gain accuracy overall. Instead, it seems that participants opted for a more conservative strategy that avoids making relatively large errors by incorrectly biasing their responses towards irrelevant priors after changepoints, and in exchange they accept a moderate precision reduction overall, i.e., increased response variance but less bias. The choice of strategy can thus be seen as an instance of the bias-variance trade-off, which has previously been proposed to underly individual differences in learning rates (Glaze et al., 2018; Eissa et al., 2022). Additionally, participants’ behavior is in line with the Ellsberg paradox (Ellsberg, 1961; Trimmer et al., 2011; Payzan-LeNestour & Bossaerts, 2011), according to which humans prefer quantifiable risk (i.e., errors with sizes defined by the experimental noise) over ambiguity (unpredictable and potentially large errors on changepoint trials).

Another factor that may have contributed to adopting a conservative strategy is the fact that observers were likely facing more uncertainty than we have included into the Bayesian models. For example, they may have been unsure about the structure of the generative process for the spatial locations of the stimuli sequences (e.g., they may have assumed that the generative means *μ*_*t*_ slowly drift through space in addition to the sudden changepoints), and they were probably not certain about the precise values of their estimates for changepoint rate and experimental noise (Payzan-LeNestour & Bossaerts, 2011). Such additional (out-of-model) uncertainty makes a conservative heuristic algorithm more attractive because the performance improvement that is gained via complex inference may only be realised when the assumed generative model is accurate (Tavoni et al., 2022). Besides merely experiencing uncertainty about the generative process, participants may have actually had a different generative model in mind than the one we used for the Bayesian model (section 4.2). For example, participants may have falsely incorporated the idea of a drifting generative mean (Nassar et al., 2010, 2019), or they may have accounted for disturbances to their prior beliefs, e.g., from memory decay (Krishnamurthy, Nassar, et al., 2017). Both examples would lead observers to undervalue the prior’s reliability and reduce their biases towards it, in accordance with the experimental data.

Lastly, diverse psychophysiological processes may have also affected the tendency to bind sensory signals with prior beliefs. For example, the arousal system has been implicated in adjusting the extent to which novel sensory information affects current beliefs (Nassar et al., 2012), as well as the extent to which prior beliefs bias perception of the sensory signals (Krishnamurthy, Nassar, et al., 2017). It has been suggested that laboratory experiments bring participants in an aroused (or sleepy) state, which therefore leads to higher (or lower) learning rates (Lee et al., 2020). In support of the hypothesis that cognitive states influence perceptual inference, it was found that a trial’s reward value, although otherwise irrelevant to task accuracy, nonetheless affected observers’ learning rates (McGuire et al., 2014).

So, there are various non-exclusive explanations for the low binding tendencies that we and others have observed. Here, we have modelled them by reducing the a-priori relevance of the priors, but there are many other modelling options that may achieve similar, or even better, fits to the empirical data. Inclusion of some of these factors into the here presented basic causal inference model should be fairly straightforward (although issues may arise with parameter identifiability), but this falls outside the scope of this study. The experimental design of the current study (Krishnamurthy, Nassar, et al., 2017) is not ideally suited to test which of the hypotheses are correct explanations for the low binding tendencies. To answer that research question, it would be useful to obtain stimulus-by-stimulus prediction responses, together with subjective judgments of uncertainty about participants’ priors, and potentially also pupillometry or neuroimaging data. However, experimental designs with overt serial decision-making risk introducing diverse cognitive biases in the response data (Talluri et al., 2018; Bévalot & Meyniel, 2023). Instead, our analysis here demonstrated the extent to which observers adhere to the core principle of Bayesian causal inference during uninterrupted sequential perception in a volatile environment.

### 3.3. Alternative models

We have derived the reduced Bayesian observer model (Nassar et al., 2012) from the fully Bayesian solution for the generative model of this changepoint task design (Adams & MacKay, 2007). In doing so, we clarified the assumptions and simplifications that underly the reduced model, and it begs the question whether different modelling choices would have led to better fits of the data.

A first consideration is that the reduced Bayesian observer model, contrary to an ideal Bayesian observer that attempts to minimize its squared errors, makes predictions about the upcoming stimulus location based on the latest posterior belief about the current generative mean. This model assumption was found to be true, as participants did not adjust their prediction responses with a bias towards the centre of space to account for the possibility of an upcoming changepoint. This finding was very clear and has been reported before (Nassar et al., 2010), so we did not consider the central bias in any of our model comparisons. But we do note that this is another indication that the fitted hazard rate parameters should not be interpreted as the expected probability of an upcoming changepoint. After all, if some participants would predict a 90% chance of a changepoint for every upcoming stimulus, then why would they not just respond near the centre of space all the time?

Second, the reduced Bayesian observer model neglects the spatial boundaries of the generative mean while it iteratively updates its beliefs. Unlike the fully Bayesian observer, it does not induce small central biases for beliefs that are near those bounds. Our model comparison analysis clearly demonstrated that incorporating the complex space truncation computations from the fully Bayesian solution made the model fits worse. This result suggests that the spatial boundaries were only accounted for when preparing to make a mouse response on the finite semicircular arc on the screen, and that the preceding computations for evidence accumulation and causal inference take place in an apparently unbounded internal representation of space. We remind the reader of the other parts of the experimental task (not analysed here) where participants had to localize a subsequently presented sound without a concurrent visual signal (Krishnamurthy, Nassar, et al., 2017). Hence, it made sense to pay particular attention to the auditory components of the stimuli, and only map the result to the visual arc on the screen when it was necessary to do so for a response. Moreover, ignoring space truncation significantly simplifies the computations at a minimal loss of accuracy, and therefore presents a logical option for an agent with limited resources (Tavoni et al., 2022; Piasini et al., 2023).

Third, we found that extending the memory capacity of the reduced Bayesian observer model did not significantly improve the fits for most of the participants. The memory limit is probably the most striking simplification in the reduced model because it deviates so much from the infinite memory requirement of a fully Bayesian observer. Yet, our analyses suggest that there is hardly any difference in the predictions of the models with various memory capacity settings as long as the weighted average pruning function (keep_*WAVG*) is used to reduce the memory load (Nassar et al., 2012). The two other pruning functions that we tested (keep_*MAXP* and keep_*RCNT*) performed objectively worse in terms of accuracy, because they explicitly discard (i.e., forget) potentially useful information (Adams & MacKay, 2007; Skerritt-Davis & Elhilali, 2018). More importantly, they also reduced the quality of the fits to participants’ responses. These results suggest that near-optimal Bayesian causal inference can be achieved with limited memory requirements by efficiently summarizing the retained prior information, and that human observers are likely to use a very similar strategy (although not nearly achieving optimality because of reasons that we discussed in the previous section).

We have chosen to restrict our analysis here to a factorial model comparison that included 42 variations of the reduced Bayesian observer model. However, naturally, other changepoint detection models have been described in the literature. For example, Wilson et al., (2013) proposed an interesting model variant for changepoint detection tasks that eludes the need for a pruning function to reduce memory load by instead attributing weights to a fixed number of nodes with predefined learning rates. Processing of sensory evidence leads to an update of the nodes’ location estimates via independent delta rules and a redistribution of the nodes’ weights via approximate causal inference. The eventual predictions are computed as a weighted average across the nodes. Because of the model’s ability to adaptively adjust the effective learning rate via the weights, its predictions will be qualitatively similar to the reduced Bayesian observer that we evaluated here. Furthermore, probabilistically weighing multiple nodes allows one to retain uncertainty about the occurrence of a changepoint, and so it resembles the behaviour of our models with memory capacities greater than one (*M* ≥ 2). However, it remains unclear how the predefined learning rates have to be determined and how they limit the model’s flexibility to fit to participants’ responses.

Remarkably, alternative modelling approaches that do not explicitly include changepoints in their generative models have also frequently been used to examine learning in changepoint environments. Notable examples are the Hierarchical Gaussian Filter (HGF; Mathys et al. 2011, 2014) and the Volatile Kalman Filter (VKF; Piray & Daw, 2020). In their generative models, volatility is implemented by allowing the latent cause of the stochastic observations to take random walks (i.e., drift) through space. Within the context of this paper, this would assume that a new generative mean is sampled from a normal distribution that is centred on the previous generative mean: 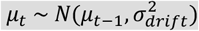. Crucially, the drift rate (i.e., variance of the random walk) is hierarchically regulated and fluctuates in time, thus producing volatile stimuli sequences. The amount of volatility over time (i.e., changes of the drift rate) is determined by a higher order parameter (similar to the hazard rate parameter in our changepoint models). During inference, the modelled observers obtain momentary estimates of the latent cause (mean and variance) and its volatility (current drift rate). Upon experiencing a changepoint, they infer a sudden jump in volatility, which leads to an increase of the learning rate for the next observation. As such, the modelled agents adapt to changepoints reasonably quickly.

However, critically, modelled predictions immediately following a changepoint (i.e., SAC 1 responses) will not be very accurate for HGF and VKF observers, because the current learning rate depends on the inferred volatility over the preceding observations, but not on the current prediction error (Eq. 51 in Mathys et al., 2011; Eqs. 9-10 in Piray & Daw, 2020). Hence, we would see a straight horizontal line when we plot these agents’ normalized biases towards the prior as a function of the latest prediction error (the line’s height would depend on the inferred volatility over preceding stimuli [averaged over multiple trials]). This model prediction is clearly at odds with the adaptive curves of the empirical data that we presented in Figure 4A. This is a fundamental difference with the method of Bayesian causal inference that we advocate for in this paper, and it is a direct consequence of the fact that the generative model of the HGF and VKF do not account for changepoints. The models adjust the prior’s reliability for the expected drift rate (i.e., our Eq. 3 is modified), but the prior’s relevance is never questioned (belief updates essentially follow Eq. 2). One possible reason for the fact that these models have nevertheless been seemingly successful at describing human decisions in changepoint environments is one that we have already put forward for fixed learning rate models: if observations are information-poor (Ryali et al., 2018), then a changepoint cannot be inferred based on a single observation (e.g., when estimating probabilities based on binary outcome variables). Only in such changepoint environments will agents that model volatility based on Gaussian random walks behave similarly to agents that account for volatility via causal inference.

### 3.4. Concluding remarks and future directions

We have shown that the reduced Bayesian observer model (Nassar et al., 2012) determines prior relevance via probabilistic causal inference and implements model averaging via its pruning function, which efficiently keeps model complexity to a minimum by retaining only a single best estimate for the source of every observation, together with a measure of uncertainty. After adjusting for individual prior binding tendencies, this simple algorithm described human predictions well in the environment for which it was designed: sequential noisy observations from static latent sources that occasionally suddenly switch location. The model would probably perform worse in a different environment that does not correspond with its generative model. However, Bayesian causal inference as a general principle is useful in any environment where there are multiple competing prior beliefs about the possible causes of observations. Accordingly, our brains seem to have adopted the method for a diverse set of tasks across the domains of perception and learning (Yu et al., 2021, Shams & Beierholm, 2022).

We have laid out the core components of Bayesian causal inference for this particular changepoint experiment (Figure 2). While we believe that the insights that we presented hold true for other changepoint tasks, the model equations will likely have to be modified to account for different environmental statistics. Specifically, we envision that the reduced Bayesian observer model forms a useful basis for modelling perceptual decisions in volatile environments that include dynamically moving sources in addition to changepoints (see also Nassar et al., 2010, 2019). It will be particularly interesting to depart from the assumption of a Gaussian random walk and instead assume that latent regular patterns underly the noisy observations. The Dynamic Regularity Extraction (DREX) model, developed by Skerritt-Davis & Elhilali (2018, 2021), is an excellent example in this direction. We took inspiration from that model for the variable memory capacity, but their arguably most intriguing contribution is that the DREX model tracks the covariance of successive observations in addition to their mean. This enables it to learn temporal dependencies in the stimuli sequences, and thus to detect deviations thereof that may indicate environmental changepoints. A drawback of the DREX model is that it is memory intensive and computationally expensive, especially when used to track covariance over time. It would be worth investigating whether the number of competing causal hypotheses (i.e., possible patterns) can be reduced by implementing some sort of weighted average pruning function, similar to the one in the reduced Bayesian observer model.

Another interesting research direction for which Bayesian causal inference lends itself naturally (more so than the classical view on belief updating expressed in terms of learning rates) is one that increases the number of active latent causes. In the changepoint paradigm, agents can simply forget their previous beliefs after a new source has been inferred for the latest observation, because the previous latent cause will not become relevant again. The opposite is true in an oddball paradigm. There, outlier observations may be caused by secondary sources that are occasionally active, but those oddball causes should be ignored, and agents are supposed to track the primary source only (Nassar et al., 2019, Bakst & McGuire, 2023). Hence, outlier signals should not be integrated with prior beliefs, but those beliefs should persist despite their irrelevance for the latest sensory input (n.b. low bias to prior *and* low learning rate!). This is akin to real world situations in which multiple objects coexist, even if temporarily obscured or silent. One can extend this line of thinking to more realistic experiments wherein causal inference is required to correctly discover latent causes of sensory signals from among multiple known objects or assign a new cause after a potential environmental change (Gershman et al., 2015). For example, a multi-source Bayesian causal inference model that is based on perceptual clustering via the Chinese Restaurant Process was recently successfully applied to explain several phenomena in auditory stream segregation (Larigaldie et al., 2024). While there is considerable psychophysical and neurophysiological evidence that supports the view that our brains use Bayesian causal inference to organize the plethora of sensory inputs according to their latent causes (Yu et al., 2021), many open questions also still remain.

## 4. Methods

The methods section is structured as follows. First, we provide details on the experimental task (section 4.1) and how the trial generation process can be translated to a generative model (section 4.2). We then derive the fully Bayesian inference solution (section 4.3), with a special emphasis on the prior relevance (section 4.4). Second, we provide details on the implementation of model variations via the pruning function (section 4.5), decision strategy (section 4.6), and late truncation simplification (section 4.7). Third, we describe the construction of a response distribution based on the inferred prediction (section 4.8), which is subsequently used to fit model parameters to participants’ responses, for various models, whose predictive performance we can then compare (section 4.9). The final two sections elaborate on the methodology that was used to compute an a-priori likelihood surface for the model parameters (section 4.10) and on specifics of the qualitative analysis for response accuracy and biases (section 4.11).

All analyses were run in Matlab 2019b (The MathWorks, Inc.) and JASP 0.19.1 (https://jasp-stats.org/). The analysis code and behavioral data are available on: https://github.com/YIRG-Dynamates/priorRelevance/.

### 4.1 Task and data

The current study is a novel analysis of the experimental data that was acquired and published by Krishnamurthy, Nassar, Sarode & Gold (2017). They made the data available to us upon request and they gave permission for us to share the relevant data for this publication in the public repository that is mentioned above.

The dataset that we received consisted of data from twenty-nine subjects who participated in between three and six experimental sessions on different days. Each session was subdivided into four main task blocks. Within each block, participants were presented with a sequence of six hundred audiovisual stimuli whose angular locations were modulated (frontal azimuth between -90° and +90°; elevation was kept constant at 0°). The sequences were paused at pseudorandom times about thirty times per block, but always after at least eight consecutive stimuli had been presented. During the pauses, participants were asked to make a prediction response about the angular location of a single sound stimulus that would be presented shortly thereafter. Participants’ pupil size was recorded in a 2.5 second delay period following the sound, after which participants made an estimation response about the sound’s location and a binary confidence judgement about that estimate. This marked the end of the pause, and the audiovisual sequence would then continue. Here, we treat each partial sequence up to the end of a pause as a separate trial, and we limit our analysis to the audiovisual stimuli and the subsequent prediction responses only, i.e., we henceforth ignore the single sound presentations and their associated pupillometry measurements, estimation responses and confidence judgments. So, for our analysis purposes, there were approximately thirty trials per main task block, each consisting of an audiovisual stimulus sequence followed by a prediction response.

The auditory and visual signals were presented in synchrony with a duration of 300 milliseconds (ms) and the interstimulus interval was 150 ms. The visual signals consisted of spatial markers on a semicircular arc that was continuously present on an isoluminant visual display in front of the participants. Responses were made on that same arc by moving a mouse cursor to the predicted location of the next stimulus. The auditory signals consisted of five 50 ms white noise pulses (incl. 5 ms on- and offset cosine ramps) that were separated by 10 ms silence. The sounds were spatialized by convolution with a head-related transfer function (HRTF), and they were presented through headphones. Each individual underwent a short test at the beginning of the first session to select an HRTF from the IRCAM database (http://recherche.ircam.fr/equipes/salles/listen/download.html) that gave the best circular experience for a sound sequence that moved through the horizontal plane in regular steps.

The spatial locations of the stimuli were sampled according to the pseudorandom generative process that is described in the next section (4.2). However, small modifications were implemented to assure that stimuli locations did not exceed the range between -90° and 90° (noise was resampled if this occurred). Furthermore, while the probability for a changepoint was in principle equal for all stimuli and the number of stimuli per trial was unpredictable, care was taken such that the last stimulus in each trial occurred equally often between 1 and 5 stimuli after the last changepoint. In other words, there was an approximately equal number of trials for stimulus-after-changepoint (SAC) levels 1 to 5. Moreover, the number of stimuli in each trial was controlled such that the average trial length (roughly twenty stimuli) was approximately equal across those five SAC levels. These mechanisms operated on a per-block basis, such that they also ensured a balanced design for two conditions with high and low levels of experimental noise that were fixed within- and alternated across blocks. The data of the first session of each participant was not analysed, i.e., the first two blocks of both noise conditions were ignored, because we assumed that participants needed this time to familiarize themselves with the task, learn about the generative structure, and estimate the changepoint probability and the amount of stochasticity in the noise conditions. Hence, between 120 and 300 trials per participant for each noise condition were included in the analyses.

### 4.2 Generative model

The pseudorandom generative process is graphically depicted in Figure 8. For each timepoint *t* a random draw from a Bernoulli distribution (*B*) determines whether or not a changepoint occurs, *cp*_*t*_ = 1 or *cp*_*t*_ = 0. The beginning of each trial always starts with a changepoint. At all other timepoints there is a constant hazard rate that determines the probability of a changepoint, *H*_*cp*_ = 0.15. Hence, *H*_1_ = 1 and *H*_*t*_ = *H*_*cp*_ for all *t* > 1. On a changepoint, the generative mean *μ*_*t*_ is drawn at random from a uniform distribution on the interval [*a* = −90°, *b* = 90°]. If no changepoint occurs, then the generative mean remains the same as on the previous timepoint, *μ*_*t*_ = *μ*_*t*−1_. Finally, stimulus location *x*_*t*_ is drawn at random from a normal distribution with mean *μ*_*t*_ and variance 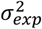, which depends on the experimental condition: *σ*_*exp*_ = 10° and *σ*_*exp*_ = 20° for the low and high noise conditions, respectively.

**Figure 8.**
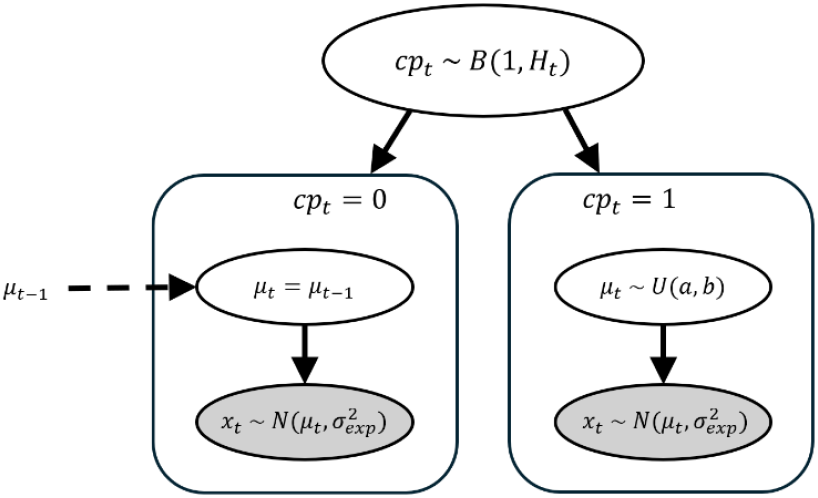
**The generative model describes the process that gives rise to sensory signals. For every stimulus, a random draw from a Bernoulli distribution determines whether or not a changepoint occurs. For a changepoint (right panel), the generative mean is drawn at random from a uniform distribution, whereas it remains the same as before for no changepoints (left panel). Finally, in both cases, the stimulus location is drawn at random from a normal distribution that is centred on the generative mean. See text in section 4.2 for details and generative parameter settings.**

### 4.3 Bayesian inference

The task is to predict the location of the next (sound) stimulus, *x*_*t*+1_, whenever the sequence stops. In order to do so accurately, an observer would be wise to continuously estimate the unknown generative mean *μ*_*t*_, because the next stimulus is likely to be sampled at random from a normal distribution that is centred on *μ*_*t*_. Hence, we will describe an algorithm that infers the location of the unknown *μ*_*t*_ from the observations *x*_1:*t*_.

While algorithms have been developed to simultaneously estimate the hazard rate, *H*_*cp*_, and experimental noise, *σ*_*exp*_, from the observations during ongoing sequence presentations (Wilson et al., 2010; Piray & Daw, 2021), we here assume that participants have learned the values for these parameters and we treat their estimates as stable constants that we denote as 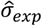 and 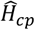.

Bayesian inference for changepoint problems of the like that we face here was described by Adams & MacKay, 2007 (also see Fearnhead and Liu, 2007). The optimal solution requires an agent to have extensive memory and computational capacity because its posterior distribution at any timepoint *t* is a weighted mixture distribution that consists of *t* nodes (i.e., mixture components): a new node is added with every new observation. Since this would quickly exhaust human working memory limits and become computationally intractable for longer sequences, we will introduce a maximum memory capacity (*M*) for the number of nodes that can be memorized and subsequently utilized. In our implementation here, this means that the posterior belief about the former generative mean, *μ*_*t*−1_, based on all preceding stimuli, *x*_1:(*t*−1)_, is first simplified to comply with the memory constraint via the function *fMC* (which will be discussed in more detail in section 4.5), before it is used as a prior distribution for the upcoming generative mean, *μ*_*t*_, conditional on the assumption of no-changepoint (i.e., *μ*_*t*_ = *μ*_*t*−1_). The memory constraint function *f*_*MC*_ ensures that the simplified conditional prior distribution consists of a weighted mixture distribution of maximally *M* nodes:

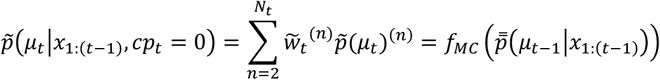

Please note that we will consistently refer to the agent’s subjective prior probabilities, distributions and parameters by means of a tilde, while we use a double overbar for subjective posterior probabilities, distributions and parameters. These are likely to be approximations of the priors and posteriors of an ideal observer because of simplifications and estimations. We will index the prior nodes at timepoint *t* with *n* = 1: *N*_*t*_, where *N*_*t*_ is defined by the sequence length (*t*) or by *M* + 1, whichever is smaller: *N*_*t*_ = min (*t, M* + 1). Node indices are superscripted within parentheses. The first prior node is reserved for the assumption of an upcoming changepoint: 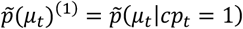. Note that given a changepoint, *μ*_*t*_ is conditionally independent of *x*_1:(*t*−1)_. The full prior distribution for *μ*_*t*_ is described as a weighted sum of the two options, either there was a changepoint or there was not:

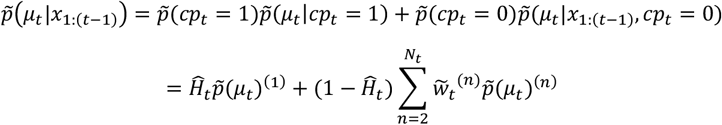

where the estimate of the hazard rate 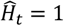 for *t* = 1, and 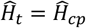 otherwise.

Upon observing *x*_*t*_, the posterior distribution for *μ*_*t*_ is computed via Bayes’ rule by multiplication of the prior with the subjective likelihood function, 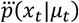, which is equal to 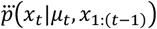 due to conditional independence, and by division with a normalization constant, *c*(*x*_*t*_):

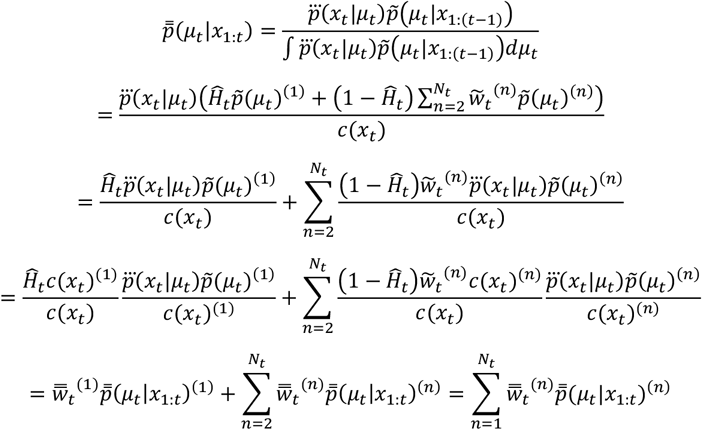

We thus find that the posterior is a weighted mixture distribution, where each of the posterior nodes is computed by multiplication of the likelihood function with the respective prior node, followed by normalization:

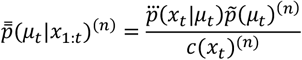

where the node-specific normalization constants *c*(*x*_*t*_)^(*n*)^ ensure that the posterior nodes are themselves proper probability distributions:

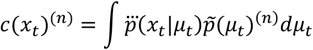

Furthermore, we have seen that the posterior weights depend on the prior weights as:

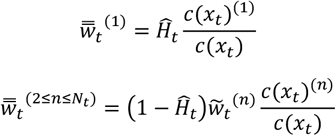

where we note that the overall normalization constant *c*(*x*_*t*_) is a weighted sum of the node-specific normalization constants:

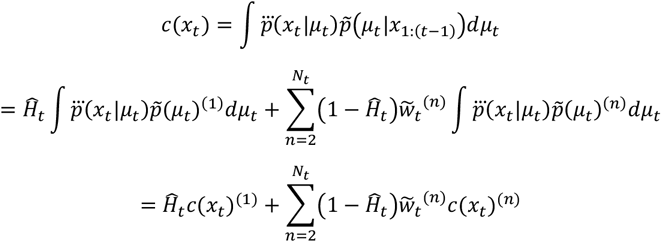

Now, we will make this general approach for Bayesian inference more concrete by showing that it can be implemented via simple updating equations for the summary statistics (i.e., parameters) of each node’s distribution.

We will start with a single observation, *x*_*t*_. Since the agent has learned that the stimulus was sampled from a normal distribution that is centred at *μ*_*t*_ with variance 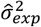, it is straightforward to construct the subjective likelihood function for *μ*_*t*_ using the symmetry of the normal distribution:

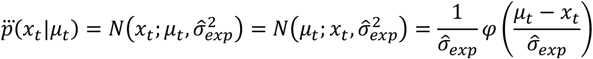

where *φ* represents the standard normal distribution.

In the case of a changepoint, *μ*_*t*_ is sampled from a uniform distribution on the interval [a, b]. Hence, the first node of the prior distribution equals:

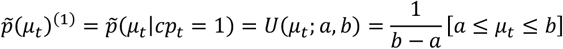

where we have used the Iverson bracket notation.

It follows that the first posterior node is a truncated normal distribution:

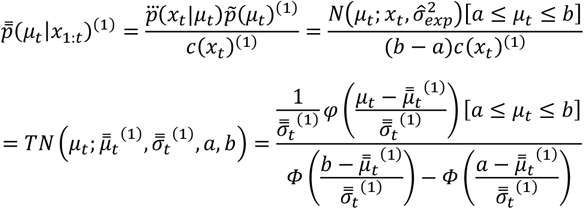

characterized by the following parameters (together with constants *a* and *b*):

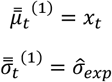

where *Φ* represents the cumulative distribution function of the standard normal distribution.

Since the first posterior node becomes the second prior node (see section 4.5 for details), the second posterior node will also be a truncated normal distribution. In fact, this is the case for each posterior node with index *n* > 1:

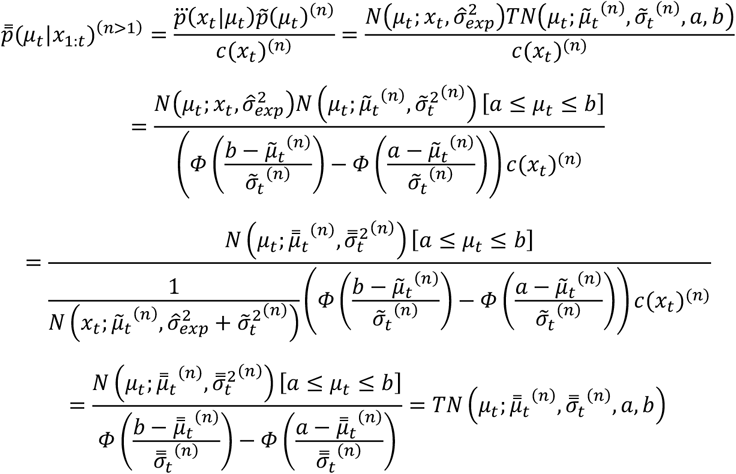

where the posterior parameters result from the product of the two normal distributions in the numerator, which itself is a scaled normal distribution. The posterior parameters can be computed efficiently with the following update equations:

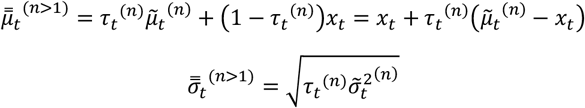

where the “prior reliability” *τ*_*t*_ ^(*n*)^ represents a normalized measure of precision (i.e., reciprocal of variance) of the untruncated prior node relative to the precision of a single observation with regards to the measure of interest *μ*_*t*_:

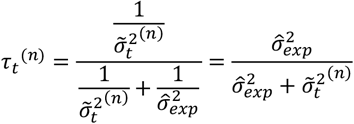

Computation of the posterior node via the normalized product of likelihood and prior node can thus be viewed as an example of precision-weighted integration.

Finally, we still need to define the nodes’ normalization constants, *c*(*x*_*t*_)^(*n*)^, which are required to compute the posterior weights 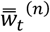:

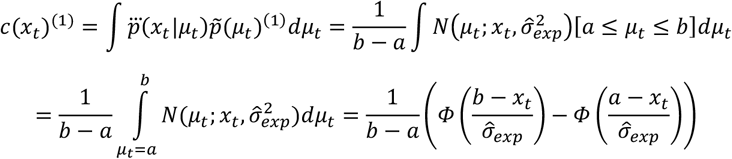

Likewise, we also integrate out the unknown *μ*_*t*_ for the nodes’ constants with indices *n* > 1:

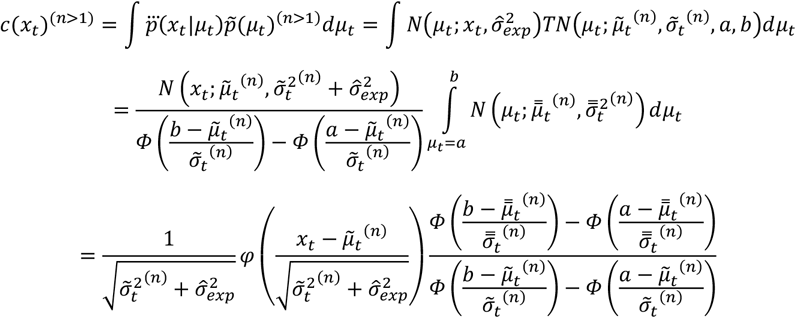

We note that these normalization constants, *c*(*x*_*t*_)^(*n*)^, can be interpreted as the likelihood of a particular node, given the latest observation *x*_*t*_. The likelihood of the first node is essentially a scalar that depends on the spatial range of *μ*_*t*_, multiplied by a term that depends on how near the observation was to the space boundaries (a and b). The likelihood of all other nodes is given by the probability density of the normal distribution that is centred on the node’s prior location parameter 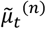, and with variance equal to the summed variances of prior and experimental noise, multiplied by another complex term that depends on the proximity of observation *x*_*t*_ and prior mean 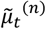 to the space boundaries.

### 4.4 Prior relevance

Since updating from prior to posterior happens on a per-node basis, and since the posterior weights, 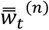, collectively form a posterior probability distribution over the nodes, we can also view these weights as a quantification of the adequacy of the respective prior nodes in terms of having provided information about the latest generative mean *μ*_*t*_. In other words, 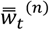 is a posterior measure for the relevance of the n^th^ prior node with regards to the latest observation *x*_*t*_, and relative to the other prior nodes.

Although the first prior node also exists, 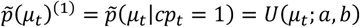, we may prefer to think of “the prior” as the collection of nodes that are based on the posterior over the previous observations, conditional on no changepoint: 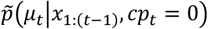. As such, we define “prior relevance” *Π*_*t*_ as the sum of the posterior weights with indices *n* > 1 (Krishnamurthy, Nassar, et al., 2017):

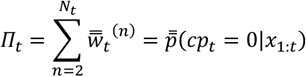

and the posterior probability of a changepoint at timepoint *t* is 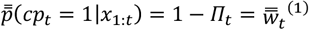.

Next, we will show that a logit transformation of the prior relevance, the unbounded quantity *Q*_*t*_, can be intuitively computed via the information-theoretic measure of surprisal:

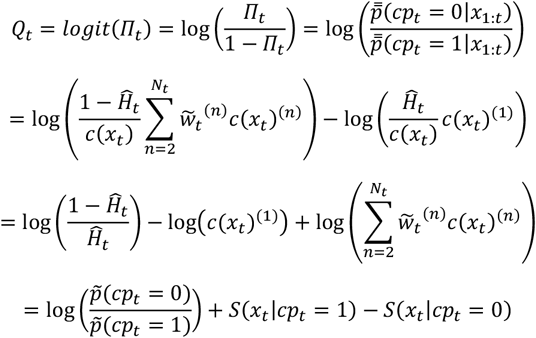

where we have defined surprisal conditional on the changepoint assumption:

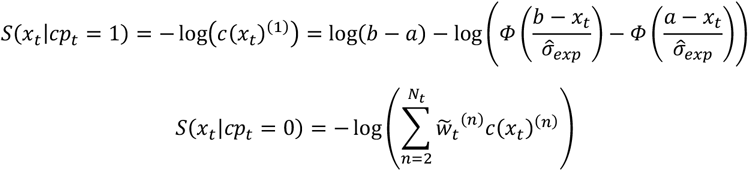

In other words, the posterior log-odds for the no changepoint hypothesis is equal to its prior log-odds plus the difference in conditional surprisal (see also Liakoni et al., 2021; Modirshanechi et al., 2022). N.b. this unbounded quantity can be transformed to a probability via the logistic function: *Π*_*t*_ = *logistic*(*Q*_*t*_) = (1 + *e*^−*Qt*^)^−1^.

Note that surprisal under the assumption of a changepoint is approximately constant, especially when the stimulus *x*_*t*_ is located in the middle of space and far away from the boundaries (a and b). Furthermore, when the prior conditional on no changepoint, 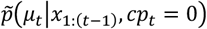, consists of only one node (*N*_*t*_ = 2, e.g., when *M* = 1), then we find that the associated conditional surprisal is dominated by the precision-weighted prediction error, i.e., the ratio of the observed squared prediction error, 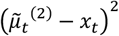, to the expected squared prediction error, 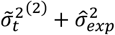:

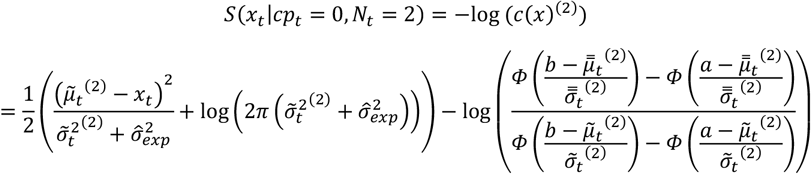

Hence, for the limited memory model with *M* = 1, the transformed prior relevance *Q*_*t*_ is equal to:

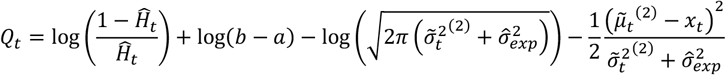

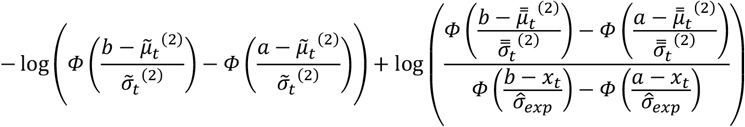

So, under the minimal memory assumption (*M* = 1), we find that the prior relevance is larger when 1) the a-priori probability for a changepoint is lower and when 2) the spatial range for the generative mean is larger. Furthermore, 3) the prior relevance decreases a-priori when the prior’s uncertainty about the upcoming stimulus location *x*_*t*_ is larger, either due to a less reliable prior on *μ*_*t*_ or due to larger experimental noise. Importantly, 4) the prior relevance decreases a-posteriori when the squared prediction error is larger, and this dependence on prediction error is stronger when the prior is more reliable and when there is less experimental noise. Finally, 5) the prior relevance increases a-priori when the prior’s location parameter is nearer to or even outside the spatial boundary for the generative mean, and 6) it increases a-posteriori when the observation *x*_*t*_ is more peripheral than the posterior’s location parameter. In other words, the prior is considered more relevant a-posteriori when it biases the percept towards the centre of space, especially when the new observation is near or outside the spatial boundaries for the generative mean.

### 4.5 Pruning function

In the above-described recursive Bayesian inference algorithm (section 4.3) we have mentioned that the preceding posterior distribution becomes the next prior distribution, conditional on no changepoint, and constrained by limited memory via the function *f*_*MC*_, for all *t* > 1:

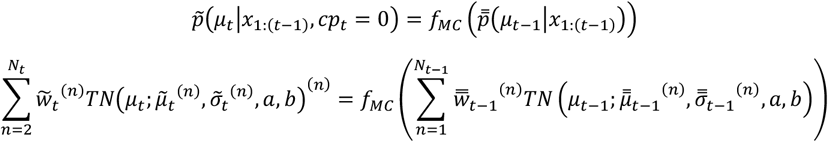

where the total number of nodes (including the first changepoint node) is: *N*_*t*_ = min (*t, M* + 1).

A fully-Bayesian ideal observer would have no memory restrictions (*M* = ∞), such that the number of nodes always increases by 1 per timestep: *N*_*t*_ = *N*_*t*−1_ + 1. However, this is unrealistic for human working memory. So, we investigated modifications of the statistically optimal algorithm.

To comply with memory constraints, it has been proposed to reduce the number of posterior nodes that carry over into the subsequent prior. This has been termed “node pruning”. Roughly speaking, it can be done in two ways: 1) one can discard posterior nodes entirely or 2) one can merge posterior nodes before prior formation. For example, 1a) Adams & MacKay, 2007 suggested to discard posterior nodes when their inferred relevance, 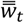, is smaller than some threshold. Instead, 1b) Skerritt-Davis & Elhilali, 2018 suggested to discard the oldest node, i.e., the one with the highest node index, whenever memory capacity is exceeded, because humans likely first forget the contribution of stimuli that were presented the longest time ago. Besides, those old stimuli are a-priori less likely to be informative for estimating the current generative mean *μ*_*t*_ in an environment with changepoints. On the other hand, 2a) Wilson et al., 2010 suggested to merge nodes based on their similarity in terms of mean and variance. Their key insight was that nodes with similar run lengths represent similar belief distributions. In our description of the Bayesian algorithm, this translates to merging nodes that have adjacent node indices. Finally, 2b) a rather extreme form of node merging was first suggested by Nassar et al., 2010, in which they reduced the posterior to one node (*M* = 1) with weighted average mean and variance, where the weights were naturally given by the nodes’ relevance, 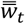. This algorithm was later refined by Nassar et al., 2012, such that the variance of the merged node was computed based on the full variance of the weighted mixture distribution, which incorporates additional uncertainty due to disparity between the means of the nodes.

Here, we incorporate some of the above insights and test which of three competitive ideas for node pruning best fits the behavioral data for various settings of the memory capacity parameter *M*: ranging from 1 to 4. In all three of the pruning methods, we make use of the insights that the oldest two nodes are often most similar, and that the memory of the oldest contributing stimuli may become hazy. So, we either merge the oldest two posterior nodes or we simply discard one of them, but we always sum their weights, so that the oldest remaining node, i.e., with the highest prior node index (*N*_*t*_) represents the belief that a changepoint occurred at least *N*_*t*_ − 1 timepoints ago. The pruning rule thus only affects the oldest pair of posterior nodes, whenever the number of posterior nodes exceeds the memory capacity, *N*_*t*−1_ > *M*. Hence, all other prior nodes are simply identical to the preceding posterior node with one index lower:

For all three *f*_*MC*_ and all prior nodes with indices 2 ≤ *n* ≤ *M*, and for prior node *n* = *N*_*t*_ if *N*_*t*−1_ ≤ *M*:

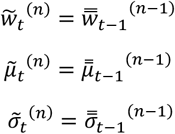

In the first pruning method we discard the oldest posterior node once the memory capacity is exceeded, in accordance with the idea of simply forgetting stimuli that were presented a long time ago (Skerritt-Davis & Elhilali, 2018). Since, we keep the most recent of the two oldest nodes, we refer to this pruning method as *keep_RCNT*:

For *f*_*MC*_*RCNT*_ and prior node *n* = *N*_*t*_ if *N*_*t*−1_ > *M*:

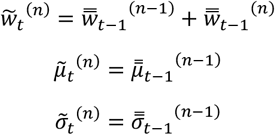

*f*_*MC*_*RCNT*_ effectively puts a limit on the number of stimuli that can get integrated, even if there are no changepoints for a long time. Hence, the variance parameter of the last prior node is bound to a minimum: 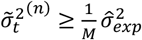, and its prior reliability is bound to a maximum: 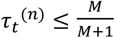

Furthermore, if the memory capacity is set to a minimum of one node, *M* = 1, this observer would only remember the location of the latest stimulus, *x*_*t*_. In this special case, *f*_*MC*_*RCNT*_ does not allow for any integration of evidence over multiple stimuli. The prior belief about *μ*_*t*_, conditional on no changepoint, 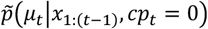 would be described by a single truncated normal distribution, 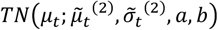, with parameters equal to: 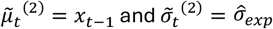.

In the second pruning method we instead discard the least relevant of the two oldest posterior nodes, i.e., we keep the node with the maximum posterior probability, *keep_MAXP*:

For *f*_*MC*_*MAXP*_ and prior node *n* = *N*_*t*_ if *N*_*t*−1_ > *M*:

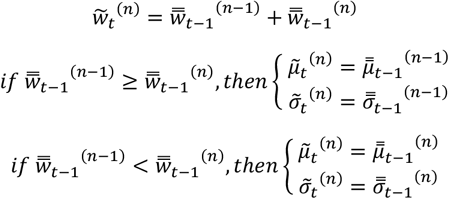

Potentially keeping the oldest but more relevant node allows one to make full use of evidence integration over many relevant stimuli (i.e., more than *M* stimuli can be integrated). In the special case with *M* = 1, an observer with the *f*_*MC*_*MAXP*_ pruning function would determine whether or not a changepoint has occurred after every stimulus (based on whether the prior relevance *Π*_*t*_ is smaller than 0.5). This observer would then keep the node that is associated with the inferred event (changepoint or not) and discard the other.

The third pruning function does not deterministically select one or the other node, but instead attempts to summarize the information from both nodes into a single one by computing a weighted average, *keep_WAVG*.

For *f*_*MC*_*WAVG*_ and prior node *n* = *N*_*t*_ if *N*_*t*−1_ > *M*:

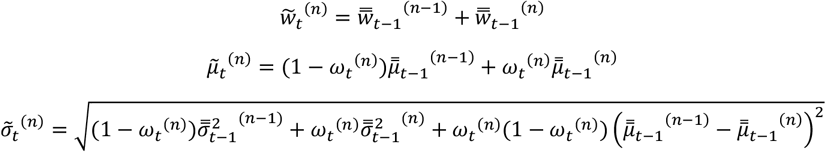

where the weight is fined as:

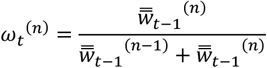

The location parameter of the new prior node is thus a weighted average of the two posterior location parameters. The squared scale parameter of the prior node is formed by a weighted average of the posterior squared scale parameters, plus a term that depends on the squared distance between the nodes’ mean parameters multiplied by a scalar that indicates the uncertainty about which of the two nodes is more relevant, *ω*_*t*_ ^(*n*)^(1 − *ω*_*t*_ ^(*n*)^), i.e., the variance of a Bernoulli distribution with parameter *ω*_*t*_ ^(*n*)^.

In the special case where *M* = 1, the *f*_*MC*_*WAVG*_ pruning function forms the core belief updating algorithm of the reduced Bayesian inference model that was described by Nassar et al., 2012. Under those minimal memory conditions, the new prior node’s parameters can be computed directly from the preceding prior node parameters and the latest observation *x*_*t*−1_ (without explicitly computing the posterior parameters):

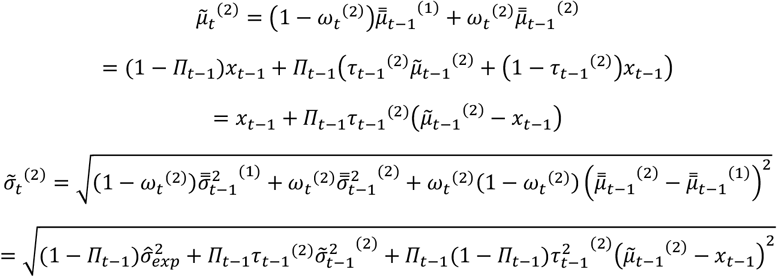

We thus find that the new prior’s location parameter is based on the latest observation plus a bias towards the preceding prior’s location parameter. The product of prior relevance and prior reliability, *Π*_*t*−1_ *τ*_*t*−1_, here serves as a normalized bias measure from the latest stimulus location (*x*_*t*−1_ at 0 in normalized space) to the prior’s location parameter (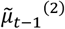 at 1 in normalized space).

### 4.6 Decision strategy

So far, we have shown how an observer can update beliefs about the generative mean *μ*_*t*_ after observing a new stimulus and how to reduce memory load by restrictive formation of a new prior. This prior distribution can then be used to predict the location of the upcoming stimulus when the observer is prompted to do so. It is straightforward to construct a predictive distribution for the unobserved stimulus *x*_*t*_ based on the prior on *μ*_*t*_ by additionally accounting for the experimental noise, 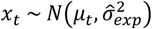:

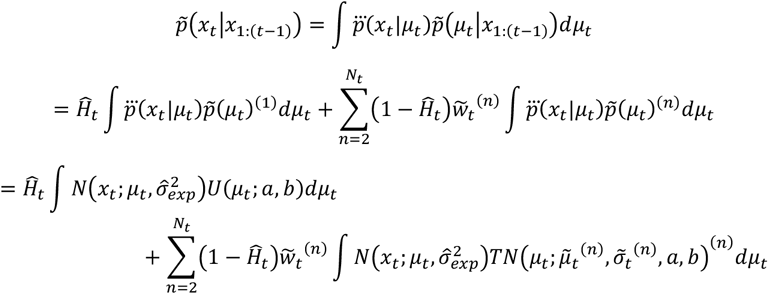

One may recognize that we have already evaluated these integrals for a particular value of *x*_*t*_ when we computed the normalization constants of the posterior nodes, *c*(*x*_*t*_)^(1)^ and *c*(*x*_*t*_)^(*n*>1)^. Fortunately, it is not necessary to evaluate the integrals for all possible values of the unknown upcoming observation *x*_*t*_. Instead, in order to make the best possible prediction, an observer only needs to compute the expected values of the predictive distribution’s nodes:

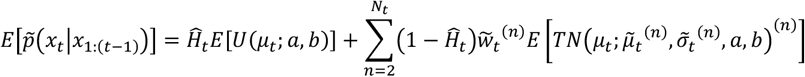

where we have used the fact that the additional experimental noise, which is sampled at random from a normal distribution, does not change the expected values of the predictive distribution’s components. These expected values are:

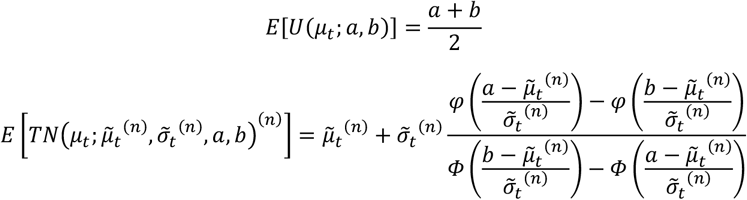

Note that the latter expectation is given by the location parameter of the prior node plus a truncation-dependent term that biases the expectation towards the centre of space, especially when the node’s location parameter 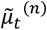 is near or outside the space boundaries for *μ*_*t*_ (a and b).

So, the statistically optimal prediction for the location of the upcoming stimulus, 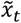, i.e., one that minimizes the squared error 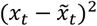 across many such predictions, is given by a weighted average of the prior nodes’ distribution expectations. Such a strategy is called model averaging (Wozny et al., 2010); where the nodes here represent competing models of the world, i.e., hypotheses about the occurrence of the latest changepoint. Since the expectation of the first prior node, conditional on an upcoming changepoint, is equal to 0.5(*a* + *b*), the ideal Bayesian observer would bias all of their responses towards the centre of space with a weight equal to 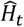. However, in accordance with Nassar et al., 2010, we did not observe such central biases in the current dataset (Figure 3A). Therefore, we assume that participants made their predictions conditional on the assumption that there will not be a changepoint. Under the “model averaging” decision strategy this amounts to:

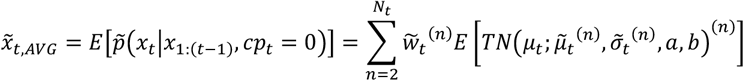

While the model averaging decision strategy is the one that a Bayesian observer would use under a squared error loss function, it is possible that participants relied on one of many other loss functions. For example, in multiple-choice tasks, the optimal strategy is to select the option with the maximum a-posteriori probability (MAP). Participants could have post-hoc applied this common strategy to determine which of the prior nodes contains the most relevant information about the generative mean *μ*_*t*_. Under the so-called “model selection” decision strategy, participants would then single out the prior node with the highest weight 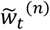 as the basis for their prediction:

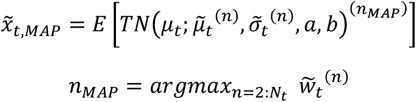

Note that both decision strategies lead to the same prediction, 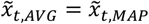, if the memory capacity is set to *M* = 1.

### 4.7 Late truncation simplification

The above near-Bayesian model computations can be simplified by ignoring the complex space truncation terms until a prediction response has to be made. In other words, during sequence presentation, sensory evidence may be accumulated (or not, in case of inferred changepoints) without considering the space boundaries for the generative mean. Under this assumption, the normalization constants (i.e., the likelihoods of the nodes) are approximated by:

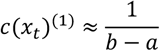

and

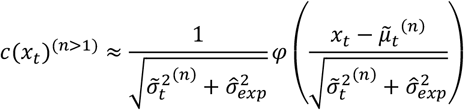

The simplification implies that any new observation *x*_*t*_ is not immediately compared against the spatial boundaries for the generative mean (a, b) when computing the prior relevance *Π*_*t*_. The spatial range merely plays a role as an a-priori offset for the transformed prior relevance *Q*_*t*_.

Nevertheless, we do assume that participants are aware of the spatial boundaries for the generative mean and that they take these into account when they make predictions for the upcoming stimulus. However, instead of computing the complex bias term for the expectation of the posterior truncated normal distributions, the prediction itself is simply truncated unto the interval of the generative mean. Hence, the predictions are first approximated:

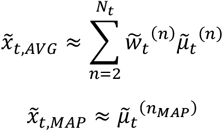

and subsequently truncated:

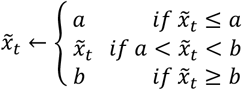

### 4.8 Response distribution

We assume that a participant intends to direct the response to the location of their best guess for the upcoming stimulus, 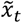, but that the response itself is inaccurate to some extent. Hence, we assume that the actual response is disturbed by random response noise according to:

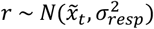

Furthermore, subjects’ attention may occasionally lapse, or they may blink during the last stimulus presentation, such that they have no idea where the next stimulus is going to be presented. On such lapses (Λ = 1), we assume that their intended response location is randomly chosen from the uniform distribution of the generative mean, 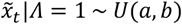. The predicted response distribution is a weighted sum of the conditional response distributions for a lapse and no lapse:

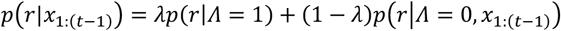

where *λ* is the lapse rate.

Since responses are bound to the response interval [*c* = −90°, *d* = 90°], we define the predicted response distribution, conditional on no lapse as a truncated normal distribution, where the location parameter is given by the model’s prediction 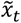:

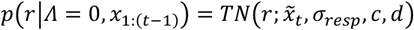

The predicted response distribution conditional on a lapse is defined by the following convolution of a uniform and normal distribution, 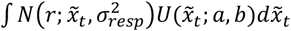, truncated to the response interval [*c, d*]:

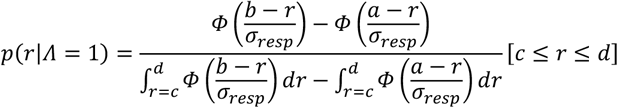

where the integrals in the denominator can be computed analytically as:

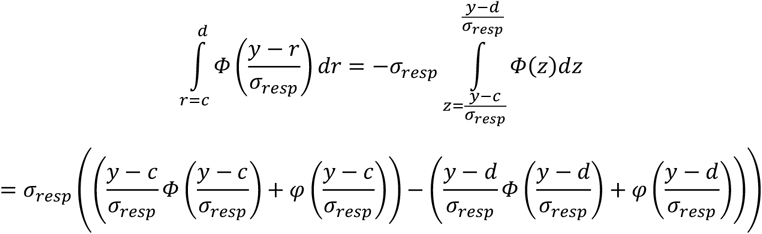

and we note that the denominator is approximately equal to *d* − *c*, such that the probability, conditional on a lapse, for any response that is well away from and in-between the space boundaries, *a* ≪ *r* ≪ *b*, is approximately constant: *p*(*r*|Λ = 1) ≈ (*d* − *c*)^−1^.

### 4.9 Model fitting and model comparison

We utilize the method of maximum likelihood estimation to optimize the parameter values of a model, *θ*_*m*_, to best predict the spatial prediction responses *r*_*k*_ of a participant, across all relevant trials *k* = 1: *N*_*k*_.

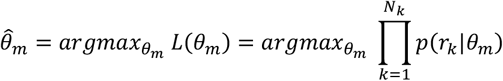

where the likelihood across trials *L*(*θ*_*m*_) for a particular set of parameter values is computed as a product of the likelihoods for each trial, and the likelihood for a single trial response is given as the probability density of the model-predicted response distribution, *p*(*r*|*x*_1:(*t*−1)_, *θ*_*m*_), evaluated at the participant’s response location, *r*_*k*_.

We fitted the models separately for both experimental noise conditions. For each fit, we optimized the following four parameters: subjective estimate of the changepoint hazard rate 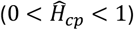, subjective estimate of the experimental noise standard deviation 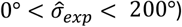, the standard deviation of the response noise (0° < σ_*resp*_ < 200°), and the lapse rate (0 < *λ* < 1):

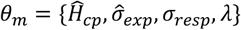

The only exceptions to this were the models with limited memory capacity, *M* = 1, and the *f*_*MC*_*LAST*_ pruning method. These models essentially base their predicted response distributions on the last stimulus location only, and they do not make use of the hazard rate estimate 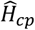. Under the late truncation simplification assumption, the experimental noise estimate 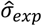 is also not used (without the simplification assumption it is used to bias the prediction 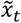 towards the centre of space via the expectation of the truncated normal). So, for these models we optimized either three 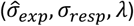 or two parameters (σ_*resp*_, *λ*) only.

To find the parameter values that maximize the likelihood we used the Bayesian Adaptive Direct Search optimization algorithm (BADS; Acerbi & Ma, 2017; https://github.com/acerbilab/bads). For each model and dataset (responses from one participant for one experimental noise condition) we ran BADS four times, each from a different starting point, *θ*_*m*,0_. To obtain promising starting points for each dataset, we first selected 1000 sets of parameter values *θ*_*m*_ at random from within the following plausible bounds: 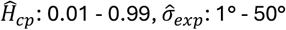, σ_*resp*_: 1° - 50°, and *λ*: 0.001 - 0.25. For each of the parameter sets we computed the likelihood *L*(*θ*_*m*_), and we subsequently selected the four parameter sets with the highest likelihood as starting points for BADS.

Non-extensive parameter recovery analyses indicated good identifiability of the true model parameters. Each of the model variations was tested several times with various parameter combinations. Responses were generated for a hypothetical observer using the experimental trials (i.e., stimuli location sequences) of a random participant from the current study as model input. Model parameters were then fitted to the generated response data using the above-described method. Approximate correspondence of the fitted parameters with the parameter values that were used during generation was checked subjectively. Although minor differences existed (as expected due to random noise generation and limited number of trials), none of the parameter recovery results gave rise to doubt identifiability.

For comparison of the models’ quality of fit to participants’ data, we computed the Bayesian information criterion (*BIC*) for each fit and then obtained an estimate of the log model evidence (*lme*) per participant for each model by summing over the two experimental noise conditions according to:

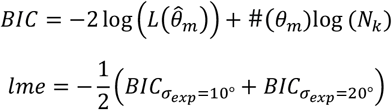

where #(*θ*_*m*_) represents the number of fitted parameters for model *m*.

The log model evidence *lme* is used to compare the models’ performances in predicting participants’ responses. We make use of two different comparison methods. First, we assume that all subjects make prediction responses according to the same underlying inference mechanism. The most likely mechanism, out of the ones tested here, is represented by the model where the summed *lme* across subjects is largest. This is known as a fixed effects model comparison. We used bootstrapping to obtain robust estimates of the summed *lme* differences between models, and their confidence intervals (Acerbi et al., 2018). Precisely, we repeatedly (N = 10,000) sampled 29 subjects with replacement and computed the summed *lme* differences for each sample, for all models relative to one reference model. Subsequently, we selected the 0.05, 0.5, and 0.95 quantiles for each model from the bootstrapped summed differences.

The second model comparison method we employed is based on the random effects method (Stephan et al., 2009). In general, this analysis accounts for heterogeneity within the subject population and estimates a probability distribution across models to indicate the likelihood of any one model to have generated the responses of a randomly selected subject. We here used the random effects method in a factorial model comparison, where probability distributions were estimated across competing model components based on the summed *lme* per subject over all models that incorporated that component (Acerbi et al., 2018): e.g., the factor ‘pruning function’ had three components: keep_*RCNT*, keep_*MAXP*, and keep_*WAVG*. For each factor, we could then compare the components’ predictive performance relative to each other via the protected exceedance probability measure (Rigoux et al., 2014).

### 4.10 Experimental noise and changepoint hazard rate inference

In the previous section we have described how we fit the model’s parameters to participants’ responses, but how did participants obtain their estimates of the parameters of the generative model? A Bayesian observer would construct a joint distribution over the parameters and update it iteratively as evidence comes in. This evidence comes in the form of a likelihood, that expresses the probability of observing a certain stimulus location *x*_*t*_ given a particular set of parameter values, and conditional on all previous observations (and the model itself). Under a non-informative prior over the joint parameter space, the parameter combination that maximizes the likelihood product across all observations forms the best estimate of the parameters. This fully Bayesian inference process is rather complex, likely computationally intractable for human brains, and falls outside of the scope of this study (for related discussions and algorithms see Wilson et al., 2010 and Piray & Daw, 2021). However, we utilized the general idea of computing likelihoods for particular combinations of parameter values to obtain an insight into which parameter combinations are more or less probable to have given rise to the given stimuli locations. Hence, we can compare the fitted parameter estimates of the participants to these likelihoods, in an attempt to quantify the errors that participants made in estimating the parameter combinations.

An equivalent algorithm to maximizing the likelihood product is minimizing the summed surprisal (Friston, 2010), across all stimuli in the experimental condition (i.e., across all stimuli in all trials with the same experimental noise). Surprisal for a single stimulus is computed as:

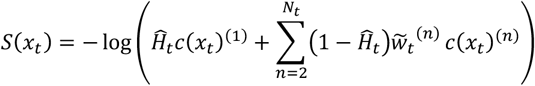

In the limited memory model (*M* = 1) with late truncation simplification, single stimulus surprisal is approximated with:

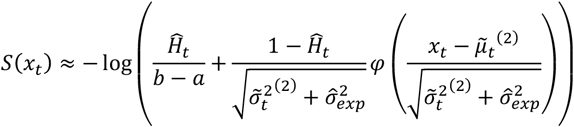

To compute the coloured background plane of Figure 5A, we computed the approximate surprisal summed across all stimuli of each experimental condition (all trials and all subjects were aggregated), for a grid of 625 parameter combinations: 25 values of 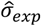 between 2.5° and 250° (logarithmically spaced) x 25 values of 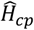 between 0.02 and 0.98 (linearly spaced). The summed surprisal values were subsequently scaled to the interval between 0 (the minimum surprisal) and 1 (surprisal for the largest 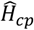 and 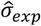, i.e., top right corner in the plot), and larger surprisal values (bottom left corner) were capped to a maximum of 1.5. Linear interpolation was used during plotting to create a smooth surface.

### 4.11 Qualitative data analysis

The data shown in Figures 3 and 4 is the group median (Q1 – Q3 in shaded area, i.e., 25% and 75% quantiles at the group level) of the median at the individual level, per grid-point on the x-axis. This robust visualization controlled for outlier responses at the individual level (e.g., lapses), and for possible outlier participants at the group level. Likewise, we employed non-parametric rank-based statistics (sign tests and Wilcoxon signed rank tests) that are robust to outliers and non-normality of the response data (which contained much intersubject variability).

In Figures 3B and 3C the trials were first discretized per SAC level (stimuli after changepoint) and the medians were computed for each bin. In Figures 3A, 4A and 4B the x-axis represented continuous variables (omniscient mean or absolute prediction error), so we instead computed rolling weighted medians for each grid point on the x-axis, where the weights were assigned to the individual’s responses by a Gaussian kernel that was centred on the grid-point (SD = 5° in Figure 3A, SD = 0.1 in Figure 4A, and SD = 0.2 in Figure 4B, see below for further explanations). To improve smoothness of the curves in the figures, the group-level medians (and Q1s, Q3s) were computed via bootstrapping (N = 100 random samples of 29 subjects, with replacement): the depicted values are the means across the bootstrapped group-level medians (and Q1s, Q3s).

The normalized errors (Figure 3C) were computed for each trial as the ratio of the participant’s error (numerator) and the error of the naïve observer (denominator), with respect to the true generative mean *μ*_*t*_:

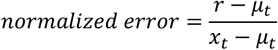

As explained in the previous paragraph, these normalized errors were discretized per SAC level and the individual’s median was then computed for each bin. The group-level medians (Q1s, Q3s) were subsequently computed via bootstrapping.

The normalized bias (Figure 4A) for each trial was computed as the ratio of the participant’s bias (numerator) and the prediction error of the omniscient observer (denominator):

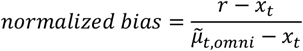

The signed bias (Figure 4B) of a trial was computed as:

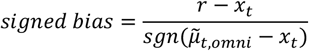

where the ‘sgn(y)’ function is -1 if y<0, and 1 otherwise.

To obtain unbiased estimates of an individual’s average trace as a function of a continuous variable (Figures 3A, 4A, 4B) via the rolling median method it is essential that the trial density across the x-axis is approximately equal. This was naturally the case for the generative mean *μ*_*t*_, because it was sampled from a uniform distribution between -90° and 90°, and so it was also approximately true for the omniscient observer’s estimate of the generative mean (Figure 3A). However, to achieve a relatively constant density of trials over the (omniscient observer’s) prediction errors for the normalized and signed bias plots (Figure 4), we used a non-linear transformation of the prediction errors when we computed the rolling medians. These transforms were based on the expected cumulative probability distributions of the prediction errors, as is explained in the following paragraphs.

The absolute distances between the generative means from before and after a changepoint are approximately distributed as a triangular distribution (across many trials):

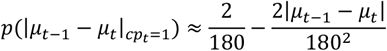

By extension, the absolute prediction errors of the omniscient observer for SAC 1 (Figure 4A) have nearly the same distribution. Hence, their cumulative distribution function (cdf) can be approximated as:

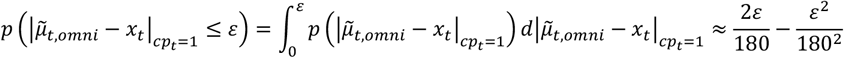

for absolute prediction errors *ε* ∈ [0°, 180°].

The cdf allowed us to assign a cumulative probability to every trial based on the omniscient observer’s prediction errors. By definition, the trials of an individual now had a constant density over these cumulative probabilities. We could thus compute an individual’s rolling median normalized bias over a grid of cumulative probabilities using a Gaussian weights kernel in the cdf-transformed space. The group-level medians (Q1s, Q3s) were subsequently obtained for each grid point via bootstrapping. Each of the grid points corresponds to a particular prediction error (of the omniscient observer), which we computed via the inverse cdf and utilized as x-axis labels in Figure 4A.

We used a similar cdf-based transformation for the prediction errors of the omniscient observer on no-changepoint stimuli (i.e., SAC 2 and SAC 3 in Figure 4B). The directional prediction errors are roughly distributed as a normal distribution:

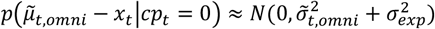

Since the omniscient observer is fully aware of the changepoints, the a-priori uncertainty about the generative mean, 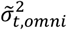, is equal to the experimental noise variance divided by the number of stimuli that were observed since the last changepoint, e.g., N = 1 for SAC 2 and N = 2 for SAC 3. Hence, the cdf for the absolute prediction errors can be written as:

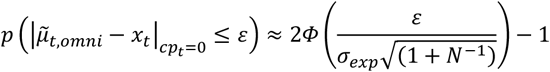

For *ε* ≥ 0, where *N* is defined by the SAC level, and *Φ* denotes the cdf of the standard normal distribution.

As before, the cdf was used to assign cumulative probabilities to trials (separately for bot noise conditions in SAC 2 and SAC 3), which were then used as a basis to compute individuals’ rolling weighted median biases on a regular grid in cdf-space. Subsequently, group-level medians (Q1s, Q3s) were computed via bootstrapping, and the corresponding absolute prediction errors of the grid points were computed via inverse cdfs. Finally, we plotted the curves of the two SAC levels on a common axis, where the axis-spacing was obtained via a cdf transformation with N = 1.5 (i.e., an average spacing for both SAC levels).

N.b. the depicted curves of the modelled observers were computed in the same way as for the human observers: i.e., we simulated their intended response locations (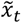, without response noise or lapses) for each individual’s set of trials and we used them to compute rolling medians at the individual level (via the cdf-transformation method, if necessary) and group-level medians via bootstrapping, with the same methodological settings as for the human observers.

## Acknowledgements

We thank Kamesh Krishnamurthy, Matthew Nassar, Shilpa Sarode, and Joshua Gold for making their experimental data available and for kindly answering our questions. We also thank Günther Koliander for helpful feedback on an earlier version of the methods section. This work was supported by the Austrian Science Fund (FWF; doi.org/10.55776/ZK66, Dynamates).

